# Nuclear RNA cap-chaperones eIF4E and NCBP2 influence distinct mRNA fates

**DOI:** 10.1101/2025.07.25.666897

**Authors:** Jean-Clement Mars, Caleb M. Embree, Biljana-Culjkovic-Kraljacic, Gokul Varma, Naiyu Shi, Aidan W.B. Carlile, Patrick Gendron, Katherine L.B. Borden

## Abstract

Gene expression is considered to converge on a single nuclear checkpoint: NCBP2 which binds the methyl-7-guanosine (m^7^G) cap on the 5’ end of mRNAs and chaperones these transcripts through all downstream processing steps as a constitutive, housekeeping activity. We show instead that a second cap-binding protein, the translation initiation factor eIF4E, also acts in the nucleus, where it and NCBP2 influence largely separate mRNA populations toward divergent outcomes rather than performing redundant, interchangeable roles. Supporting this, eIF4E and NCBP2 overexpression or genetic reduction produced distinct transcriptomic and splicing signatures affecting thousands of transcripts but only a few shared splicing events. We propose a model, mRNA Specification by Cap Chaperones (SOCCs), in which selective association of transcripts with specific cap-chaperones influences their processing. We show that altering cap-chaperone levels changes the association of specific transcripts in the nucleus with eIF4E or NCBP2 and is accompanied by altered processing outcomes that can affect the structure and function of the resulting mRNAs. Multiple properties underpin SOCCs, including spatial segregation of eIF4E and NCBP2 within the nucleus and non-overlapping cap-chaperone protein-partner networks as well as differential influence of eIF4E and NCBP2 on production of the processing machinery itself. Together, these findings suggest cap-chaperone selection as a previously unrecognized regulatory layer in gene expression, positioned to influence all nuclear mRNA processing steps as well as nuclear mRNA export.

## Introduction

Gene expression encompasses the processes involved in converting genetic information into functional protein. In eukaryotes, an essential step in this process is the addition of the methyl-7-guanosine (m^7^G) cap on the 5’ end of coding transcripts^1–3^.

Capping of mRNA is conserved in animals, plants and fungi^1–3^. m^7^G-caps are added to coding mRNAs at or near sites of transcription^1–4^. Ingrained in the central dogma of gene expression is the necessity for m^7^G-capped transcripts to be chaperoned through processing steps and subsequently escorted to the nuclear pore for export to the cytoplasm for translation^5–7^. The major purpose of the cap-chaperone is to mediate interactions between mRNAs and the appropriate processing machineries to enable maturation^6^. To date, this chaperone role was solely attributed to the nuclear cap-binding protein NCBP2 (CBP20) and its co-factor NCBP1 (CBP80) (Figure 1A, top panel). NCBP1 binds directly to NCBP2 and other RNA-binding proteins to direct mRNA fate^6,8,9^. Like the cap, NCBP2 and NCBP1 are evolutionarily conserved across plant, animal and fungi kingdoms^5,6,10^. In this model, the NCBP2/NCBP1 heterodimer (also known as the cap-binding complex, CBC) acts in mRNA processing and nuclear export, with a handover of mRNAs in the cytoplasm to the major cytoplasmic cap-binding protein eIF4E which engages transcripts in steady-state translation^11,12^.

**Figure 1:**
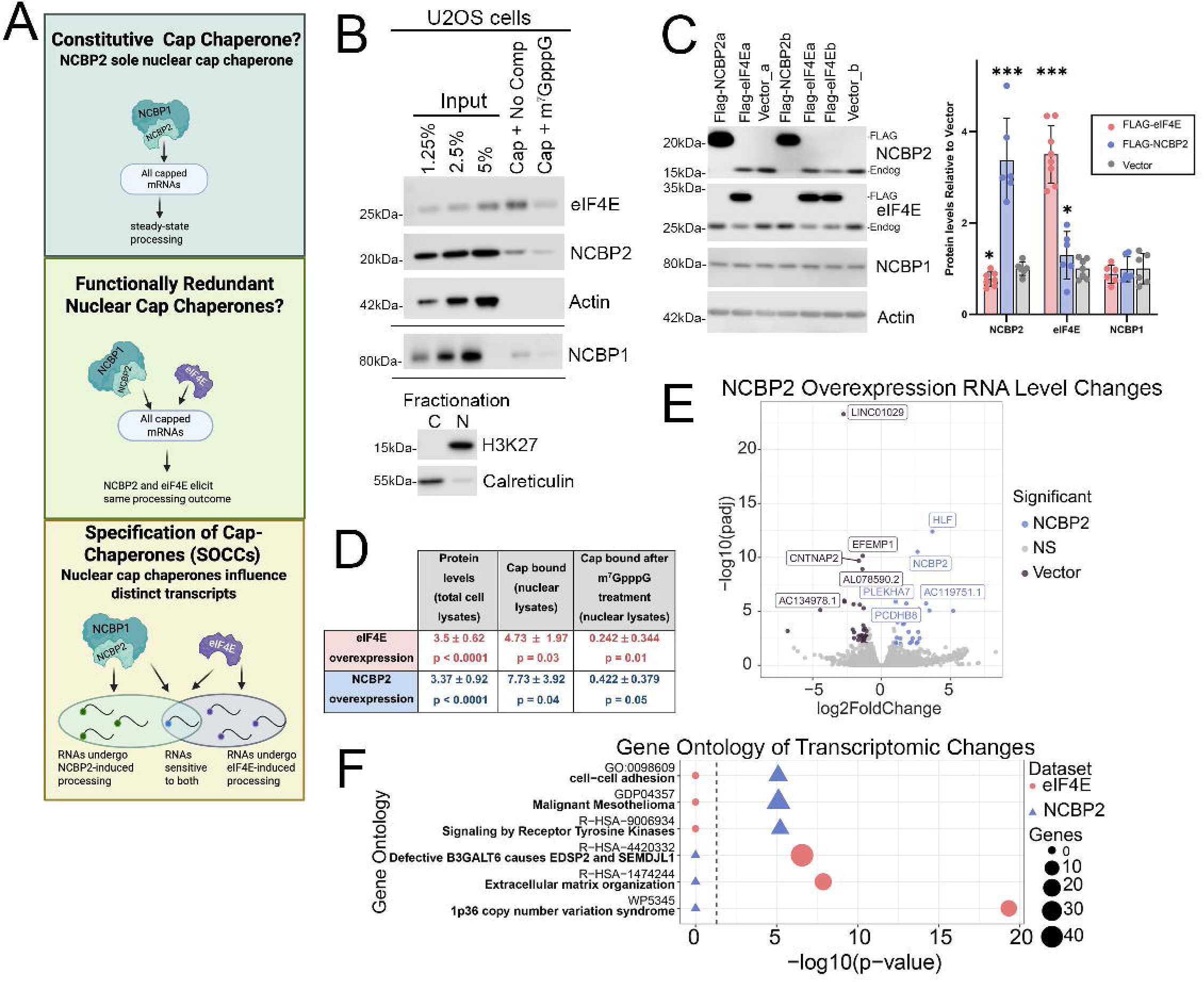
eIF4E and NCBP2 overexpression have distinct transcriptomic changes. **A)** The potential models of cap chaperones interacting with nuclear mRNA. Created with https://BioRender.com **B)** *Top* Western blot showing a representative biological replicate depicting the amount of NCBP2 and eIF4E bound to m^7^G beads treated with buffer or m^7^GpppG. *Middle* Western blot showing a representative biological replicate depicting the amount of NCBP1 bound to m^7^G beads treated with buffer or m^7^GpppG. *Bottom* Western blot of the fractionation controls for the nuclear extract used in cap chromatography. **C)** *Left* Representative western blot depicting the protein levels of eIF4E and NCBP2 upon overexpression in U2OS cells from two clones (a and b). *Right* Quantification of 6 biological replicates of NCBP2 and 7 biological replicates of eIF4E overexpression. * = p < 0.05, ** = p < 0.01, *** = p < 0.005, all others are non-significant. **D)** Western blot quantification of total protein levels and m7G-cap bead binding ± excess m7GpppG competitor. Values are shown as fold-change vs vector ± standard deviation. For total protein levels, comparisons were between the total protein content of eIF4E (red) or NCBP2 (blue) overexpression vs vector cells. For cap bound protein levels, comparisons were between the amount of eIF4E or NCBP2 bound to m7G beads in overexpression vs vector cells. p-values calculated via Welch’s t-test in PRISM. **E)** Volcano plot depicting the transcripts that are significantly upregulated (blue) or downregulated (purple) upon NCBP2 overexpression in U2OS cells. **F)** The top gene ontology terms of genes undergoing differential expression in eIF4E (red circles) or NCBP2 (blue triangles). Size of the point indicates the number of genes and position along the x-axis indicates enrichment of the GO term. Dashed line indicates p = 0.05.

Much that underpins the above model arises from the misconception that NCBP2 is the only cap-binding factor in the nucleus (Figure 1A). Indeed, eIF4E is found in the nucleus and cytoplasm in plants, animals and fungi^10,13,14^. This evolutionary conservation suggests a nuclear role. Multiple independent lines of evidence confirm that eIF4E retains cap-binding activity in the nucleus: 1) it physically associates with over 3000 nuclear mRNAs by RNA immunoprecipitation (RIP); this association is competitively disrupted by the m^7^G cap analogue m^7^GpppG but not by the control GpppG, in both RNA immunoprecipitation and gel-shift assays from nuclear lysates; 2) a cap-binding-deficient mutant (W56A) abrogates the mRNA interaction in the nucleus; and 3) the nuclear factor PML, which reduces eIF4E’s m^7^G cap affinity ∼50-fold, correspondingly reduces its nuclear mRNA association and subsequent mRNA export activity^15–18^.

Neither eIF4E nor NCBP2 requires overexpression to bind capped mRNA in the nucleus^17–20^, indicating this reflects the behavior of the endogenous proteins rather than an artifact of ectopic expression. Consistent with these observations, eIF4E and NCBP2 have comparable cap affinities *in vitro.* Purified NCBP2 alone binds cap analogues ∼470-fold more weakly than purified eIF4E (K_d_ of 84 µM vs. a median of 180 nM^21–23^); however, NCBP2’s cofactor NCBP1 induces a conformational change that increases NCBP2’s cap affinity ∼875-fold, to a median K_d_ of 96 nM^21,24^ which is within the range of eIF4E’s own affinity. Thus in the nucleus, eIF4E and NBCP2 are both positioned to bind m^7^G-capped mRNA which could have important consequences for mRNA fate.

To date, the role of eIF4E in the nucleus was largely considered to be limited to mRNA export of specific m^7^G-capped-RNAs^15,17,25–29^. Specifically, eIF4E promotes export of oncogenic mRNAs to the cytoplasm thereby increasing their cytoplasmic availability to ribosomes without changing their total levels. eIF4E physically associated with capped RNAs that contain the eIF4E-sensitivity element (4ESE) in the 3’UTR which in turn recruits co-factors including LRPPRC and XPO1/CRM1 to engage the nuclear pore complex^16,29–31^. Using a combination of molecular biology, biophysical, and NMR methods, it was shown that LRPPRC uses its C-terminus to directly bind to the 4ESE element in the target mRNA and its N-terminus to directly bind the dorsal surface of eIF4E, a region distal to its cap-binding site, while eIF4E binds the m^7^G cap^29–31^. These studies showed that the 4ESE RNA-LRPPRC-eIF4E-m^7^G cap complex is formed and then LRPPRC engages XPO1 to exit the nuclear pore complex^30,31^. By contrast, NCBP1/NCBP2 is involved in bulk mRNA export binding NXF1/NXT1 but not LRPPRC or capped-LacZ-4ESE mRNA in the nucleoplasmic fraction^31^. This provides an example of selection of one cap-chaperone over the other to engage the export of specific capped mRNAs based on co-factors and mRNA USER codes.

Consistent with a more general role in capped mRNA metabolism, recent studies revealed that eIF4E, similar to NCBP2/NCBP1, physically interacted with the capping, cleavage and polyadenylation, and splicing machineries as well as some unspliced mRNAs and their corresponding spliced products in nuclear lysates^17,26,27,29,32,33^.

Interestingly in the nucleus, the physical interaction between eIF4E and splicing factors was m^7^G cap- and RNA-dependent^33^; eIF4E physically interacted with specific capped mRNAs that contained introns and their corresponding spliced forms^33^. Similar observations establish that NCBP2 binds pre-mRNAs, mRNA processing machinery and influences processing steps including splicing^34–37^. Like eIF4E, NCBP2/NCBP1 interacts with splicing factors in a m^7^G cap-dependent and RNAse treatment sensitive manner in nuclear lysates^38^. Thus, there appears to be two cap-chaperones in the nucleus positioned to modulate mRNA fate.

Given eIF4E and NCBP2 co-exist in the nuclei of yeast, animal and plant cells^9,10,13–15,39^ and both bind pre- and mature capped mRNAs in nuclear lysates, it seems likely that there is an evolutionarily conserved purpose for this coexistence and thus it is important to understand their functional interplay. With this in mind, we asked whether the presence of both eIF4E and NCBP2 reflects functional redundancy that safeguards gene expression fidelity or is an unrecognized regulatory step in the control of gene expression (Figure 1A, middle and bottom panels respectively). We found that eIF4E and NCBP2 overexpression, and conversely their genetic knockdown, produced distinct splicing profiles with minimal overlap, arguing against simple redundancy. We showed that mRNA fate for specific transcripts could be altered by modulating the levels of eIF4E and NCBP2. We formulate a new model whereby selective association of transcripts with specific cap-chaperones influences mRNA fate, a model we refer to as mRNA Specification by Cap-Chaperones (SOCCs; Figure 1A bottom panel). Our data indicate that this may be a regulatory step in gene expression that can influence the structure of the mRNA and its final protein form. We identify molecular features that are consistent with cap-chaperone selectivity and its consequences for mRNA fate. SOCCs proposes a previously unrecognized step in the control of gene expression associated with thousands of mRNAs, representing a substantial departure from the current NCBP2 only cap-chaperone paradigm.

## Results

### eIF4E and NCBP2 elicit divergent transcriptomic and splicing signatures

We investigated whether the co-existence of nuclear eIF4E and NCBP2 reflected redundancy in processing to protect the fidelity of gene expression or if mRNAs were specified for a particular cap-chaperone protein to influence their fates. To monitor this, we used human osteosarcoma cells (U2OS) which are characterized by nuclear and cytoplasmic eIF4E and NCBP2 and by eIF4E proteins levels similar to healthy volunteers^33^. As a first step to address this, we employed cap-chromatography to confirm that both endogenous eIF4E and NCBP2 bound to the m^7^G cap in the nucleus. Nuclear lysate quality was confirmed using H3K27 and Calreticulin fractionation controls for the nuclear and cytoplasm fractions respectively. Nuclear lysates were incubated with m^7^G-agarose beads, and cap-chaperone binding assessed by western blot (Figure 1B). Addition of free m^7^GpppG reduced the association of either eIF4E or NCBP2, as well as the other component of the CBC, NCBP1, with the m^7^G-cap agarose indicating that the association of each cap-chaperone with beads was cap-specific. To extend these findings, we also used nuclear lysates from the MM6 AML cell line which is characterized by inherently elevated eIF4E levels. Consistently, we observed eIF4E, NCBP2 and its partner protein NCBP1 bound to the cap columns and binding was abrogated upon addition of excess m^7^GpppG (Figure S1A).

To study these cap-chaperones, we generated stable overexpressing FLAG-eIF4E and FLAG-NCBP2 clonal cell lines as well as parallel vector control lines. In whole cell lysates, we observed a 3.5-fold increase in eIF4E relative to vector controls by western blot (Figure 1C) which was well within the range of elevation observed in cancer patients (>3-8 fold)^33,40–42^. In whole-cell lysates, eIF4E overexpression did not substantively alter levels of endogenous NCBP1 or NCBP2 relative to vector controls (Figure 1C). Conversely, FLAG-NCBP2 levels were similarly elevated by ∼3.5 fold relative to vector controls which were also similar to those observed in cancer patients (Figure 1C)^43–45^. To ensure that overexpression led to increased levels of cap-chaperone bound to cap columns, we generated nuclear lysates and compared cap-column binding of endogenous eIF4E or NCBP2 in vector controls to levels in FLAG overexpression cells. Indeed, we observed a very close correlation between eIF4E overexpression (∼3.5-fold) and amount of eIF4E upon overexpression bound to the cap beads (∼4.7-fold; Figure 1D). NCBP2 overexpression led to ∼7.5 fold enrichment of NCBP2 in FLAG-NCBP2 cells relative to vector controls (Figure 1D). Thus, overexpression of NCBP2 and eIF4E produced active cap-binding proteins that roughly scales with the level of overexpression.

To assess whether eIF4E and NCBP2 elicited redundant or divergent outcomes to gene expression, we compared the transcriptional signatures of eIF4E and NCBP2 stable overexpressing clonal cell lines. In order to best monitor both pre-mRNA and mRNA, we employed ribodepletion for library preparation. We first examined whether eIF4E impacted steady-state transcription using our previously reported RNA-Seq data comparing stably overexpressing 2FLAG-eIF4E U2OS cells to vector controls, using 3 different clones per condition (<u>GSE158728</u>)^33^. eIF4E overexpression was not associated with global reprogramming of the transcriptome, with 402 differentially expressed genes (p_adj_<.05) in Vector versus 2FLAG-eIF4E U2OS cells of which 333 had > 2-fold change (<u>GSE158728</u>)^46^. These transcripts were approximately equally distributed between elevated and repressed in eIF4E overexpressing cells (233 and 100 respectively) out of 5738 annotated transcripts observed with >10 TPM including both coding and non-coding RNAs (<u>GSE158728</u>)^27^. Metascape analyses of these targets revealed top protein-protein interaction network enrichment included “TNFR2 non-canonical NF-κB pathway”, “Defective B3GALT6 causes EDSP2 and SEMDJL1” and “positive regulation of DNA metabolic process” (Supplemental Table 1). The enrichment of NFkB signalling is interesting given eIF4E is a known target of NFkB^47^. Whether eIF4E’s impact on transcription is directly related to its physical association with active sites of transcription as described below remains to be determined. In all, eIF4E only influences steady state levels of ∼6% of detected transcripts by >2-fold^33^ (<u>GSE158728</u>).

Next, we assessed the impacts of NCBP2 overexpression on the transcriptome. We generated stably overexpressing 2FLAG-NCBP2 and vector U2OS cell lines monitoring two NCBP2 clones at 3 different passages and 3 different vector clones, RNAs underwent ribodepletion and then we conducted RNA-Seq to monitor changes to steady-state transcripts levels and splicing and compared these with the above eIF4E data. Differential expression analysis revealed that 101 transcripts had altered steady state levels upon NCBP2 overexpression relative to vector cells (>2-fold, FDR <.05; 55 reduced and 46 increased in NCBP2 than vector) (Figure 1E, Supplemental Table 2) (<u>GSE300496</u>). This represents ∼1.2% of the total 8331 annotated transcripts detected with >10 TPM. Metascape analyses of the transcripts with altered expression levels revealed distinct enrichment patterns for eIF4E and NCBP2 overexpression. Top GO terms for NCBP2 include “Signaling by Receptor Tyrosine Kinases”, “Malignant Mesothelioma”, and “Cell-cell adhesion” while top enriched terms for eIF4E include “p36 copy number variation syndrome,” “Defective B3GALT6 causes EDSP2 and SEMDJL1” and “extracellular matrix organization” (Figure 1F and Supplemental Table 3&4). The biological pathways driven by eIF4E and NCBP2 overexpression differed although both were cancer-related (Figure 1F). A comparison of transcripts altered at steady state by eIF4E and NCBP2 revealed 3 transcripts in common: *PALM3, INHBE,* and the long non-coding RNA *MSC-AS1* (Supplemental Table 5). Thus overall, eIF4E and NCBP2 elicit distinct changes to steady state transcript levels.

To establish if NCBP2 and eIF4E produced redundant or divergent impacts on processing, we compared the splicing outcomes of NCBP2 and eIF4E overexpression. We note that did not observe induction of splice factor mutations common in cancers e.g. SF3B1, U2AF1 or SRSF2 upon eIF4E or NCBP2 overexpression^33^. We assessed the impact of NCBP2 overexpression on splicing by analyzing the above RNA-Seq data with replicate Multivariate Analysis of Transcript Splicing (rMATS)^48^ (Supplemental Table 6). rMATS calculates the “inclusion level differences” for splicing events such as exon skipping (SE), inclusion of mutually exclusive exons (MXE), intron retention (IR) and alternative splice site usage (A3’SS and A5’SS). We note that the previously reported sign for eIF4E inclusion level difference was opposite^33^ to the NCBP2 overexpression analysis reported here and was modified for this report to match the directionality of the NCBP2 data. Here, a value of +1 for SE events indicated that the relevant exon is 100% included in 2FLAG-eIF4E or 2FLAG-NCBP2 cells and 0% in Vector cells. Using rMATS, a comparison of NCBP2 relative to vector revealed that 3894 splicing events for 2524 genes were altered using the cutoffs we previously used to examine alternative splicing changes in eIF4E (FDR-adjusted p-value <0.15; Inclusion differences of >0.05 or <-0.05). Using more stringent cutoffs that we will use for subsequent analyses (FDR <0.05, Inclusion level differences of >0.1 or <-0.1), we observed for 1537 events for 1219 genes (Figure 2A, Supplemental Table 6). The RNA-Seq experiment contained a total of 60564 annotated transcripts, 8331 of which had >10 TPM in all samples meaning that ∼15-30% were differentially spliced in an NCBP2-dependent manner depending on the cutoffs used. Only two targets, *LGR5* and *SLC7A10,* were differentially expressed and underwent AS upon NCBP2 overexpression, suggesting that NCBP2-related alternative splicing (AS) is not generally correlated to mRNA levels under these conditions (Figure S1B).

**Figure 2:**
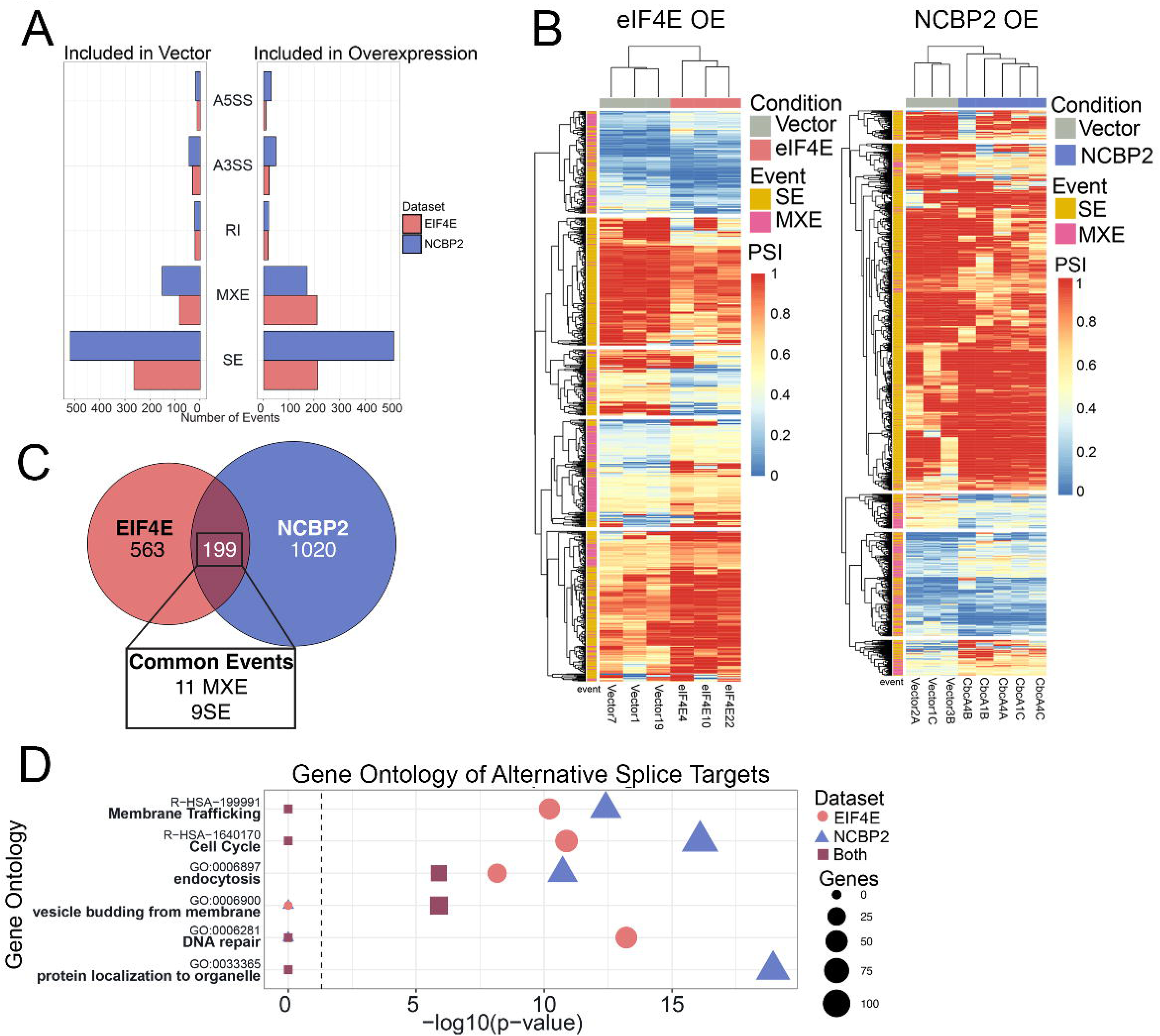
eIF4E and NCBP2 overexpression alter the splicing of distinct targets. **A)** Bar graph illustrating the number of alternative splicing events in EIF4E (red) and NCBP2 (blue). **B)** Heatmap of significantly changing SE (yellow) and MXE (pink) events showing inclusion (red) or exclusion (blue) of the exon in each sample of eIF4E overexpression (left) and NCBP2 overexpression (right). **C)** Euler plot comparing the genes that have significantly changing alternative splicing in eIF4E (red) or NCBP2 (blue) overexpression. The number of SE and MXE events that are changing in both overexpression datasets are indicated. **D)** Top gene ontology terms of genes undergoing alternative splicing in NCBP2 only (triangle, blue), EIF4E only (circle, red), or both (square, purple) overexpressions. Size of the point indicates the number of genes and position along the x-axis indicates enrichment of the GO term. Dashed line indicates p = 0.05.

In terms of splicing targets, upon NCBP2 overexpression there are ∼2.5 times as many genes than those modulated by eIF4E and ∼3.0 times as many splicing events: 886 splicing events of 762 genes in eIF4E-overexpressing U2Os (FDR-adjusted p-value <0.15; Inclusion level differences of >0.05 or <-0.05) and 548 splicing events for 487 transcripts (FDR <0.1, inclusion level differences of >0.1 or <-0.1) with 5738 annotated transcripts with >10 TPM^33^. To maintain consistency with our previous analysis^33^, we will continue to examine splicing changes caused by eIF4E overexpression using cutoffs of FDR-adjusted p-value <0.15; inclusion differences of >0.05 or <-0.05. Like NCBP2, only ∼2% (13 transcripts) had both splicing and steady-state levels altered by eIF4E^33^. This gives a total of 8-13% of transcripts undergoing AS for eIF4E, and 15-30% observed transcripts for NCBP2 depending on the cutoffs used. In all, the two cap-binding proteins elicited broad but not universal impacts on splicing signatures and effects were more restricted on transcript levels.

In terms of NCBP2, SE were by far the most frequent event (∼67%) followed by MXE events (∼21%) with an approximately equal distribution between positive and negative inclusion level differences upon NCBP2 overexpression relative to vector (Figure 2A), indicative of reprogramming of splicing for selected mRNAs rather than a global alteration to splicing. While there were some non-coding RNAs, coding mRNAs predominated (Supplemental Table 6&7). As SE and MXE events were the most common, we focused the remainder of our analyses on these events. Unsupervised hierarchical clustering analyses based on the inclusion levels across replicates revealed that these events segregated into vector and overexpression groups for both eIF4E and NCBP2 (Figure 2B). We noted no dependency of NCBP2-dependent splicing on exon length or GC content within introns (Figure S1C&D) but similar to our observations for eIF4E-overexpression, introns of alternatively spliced transcripts that were modulated by NCBP2 overexpression were ∼1.5 times or more longer than average introns (Supplemental Table 8). Consistent with previous studies in other cellular contexts^9,49^, we noted a significant bias towards the first three exons being targeted upon NCBP2-overexpression but note that other exons were also targeted (Figure S1E). A reanalysis of eIF4E data employing the same strategy also demonstrated a significant bias for eIF4E transcripts for first, second and third exons for SE events; however, like NCBP2, other exons were also impacted (Figure S1E). In all, both NCBP2-dependent and eIF4E-dependent splicing is predominated by SE events with substantial amounts of MXE events in U2OS cells (Figure 2B). Both cap-chaperone splicing programmes are characterized by a tendency to be in exons closer to the 5’end compared to all exons and involve unusually long introns; but no correlation was observed regarding GC content in these introns or length of targeted exons for either NCBP2 or eIF4E overexpression^33^ (Figure S1C-E).

Next, we established whether eIF4E and NCBP2 drove redundant splicing programmes, as would be expected for cap-chaperone housekeeping activity, or if cap-chaperones drove divergent programmes. Strikingly, a comparison of their splicing signatures revealed only 199 transcripts in common; comprising ∼16% of the genes with altered splicing induced by NCBP2 and 26% of genes with altered splicing induced by eIF4E (Figure 2C; Supplemental Table 9). An ontology analysis of genes with splicing altered by eIF4E or NCBP2 overexpression revealed strongly enriched pathways “endocytosis”, “cell cycle”, and “membrane trafficking” in common (Figure 2D; Supplemental Table 10-12). These support that while the targets may be different, there are common pathways driven by eIF4E and NCBP2. Interestingly, each group of alternate splice target genes were also involved in unique pathways; “protein localization to organelles” in genes targeted by NCBP2, “DNA repair” in genes targeted by eIF4E, and “vesicle budding from membrane” in genes affected by both proteins were highly enriched (Figure 2D, Supplemental Table 10-12). Despite the fact that the mRNA targets of eIF4E and NCBP2 share common properties, particularly a bias toward the 5’ end of exons, long introns and SE events, and are involved in some of the same biological processes, these cap-binding proteins elicit splicing and transcriptomic signatures that diverge for ∼80% of affected transcripts supporting the SOCCs model (Figure 1A, bottom panel). Consistently, the similarity in AS-dependent pathways between NCBP2 overexpression in U2OS cells and NCBP2 overexpression or knockdown in mouse cardiomyocytes^37^ (Figure S1F, 13-14% respectively), and the similar enriched GO pathways in transcripts alternatively spliced by NCBP2 overexpression in humans and mice (Supplemental Table 11&13) implies that the ability of NCBP2 to regulate AS of specific transcripts is conserved across species.

Our above assessment allowed us to identify ontological similarities and differences between eIF4E and NCBP2. Next, we established the number of overlapping splicing events to quantify redundancy between these cap-chaperones. We observed that only ∼9 SE events (∼1% of NCBP2 SE events) and 11 MXE events (∼3%) (Figure 2C, Supplemental Table 14) were conserved between eIF4E and NCBP2 overexpressing cells. Thus, while there was overlap in the transcripts affected, there were very few overlapping events between the two cap-chaperones.

### Reduction in eIF4E and NCBP2 influence distinct sets of splicing events

eIF4E and NCBP2 overexpression recapitulated cancer models in terms of divergent alternative splicing effects; next, we sought to understand whether eIF4E and NCBP2 differentially modulated splicing at baseline. We genetically reduced eIF4E and NCBP2 levels using siRNA. Knockdown was compared to the negative control siRNA to luciferase by western blot (Figure 3A). Genetic reduction led to a 3.2-fold decrease in eIF4E protein levels, and a 3.5-fold reduction in NCBP2 levels (Figure 3A). Thus, the knockdowns were comparable. Genetic reduction of NCBP2 did not alter endogenous eIF4E levels, while eIF4E knockdown modestly raised NCBP2 levels 1.3-fold. Both eIF4E and NCBP2 knockdown altered levels of NCBP1, causing a 1.5-fold elevation and 1.6-fold reduction respectively (Figure 3A). This latter finding suggests that NCBP1 is more stable as part of the NCBP1/NCBP2 heterodimer than alone in U2OS cells. As Gebhardt and colleagues demonstrated that NCBP2 knockdown did not affect NCBP1 levels in HeLa cells^19^, the influence of cellular context on NCBP1 warrants further investigation. Because NCBP1 and NCBP2 function together as the CBC, the NCBP2 knockdown phenotypes described below cannot be attributed to NCBP2 alone and are interpreted as reflecting depletion of NCBP2 accompanied by reduced NCBP1 (and thus reduced CBC); we refer to this condition as NCBP2 knockdown with concomitant NCBP1 reduction. Similarly, the 1.5-fold elevation of NCBP1 upon eIF4E knockdown should be considered when interpreting those data.

**Figure 3:**
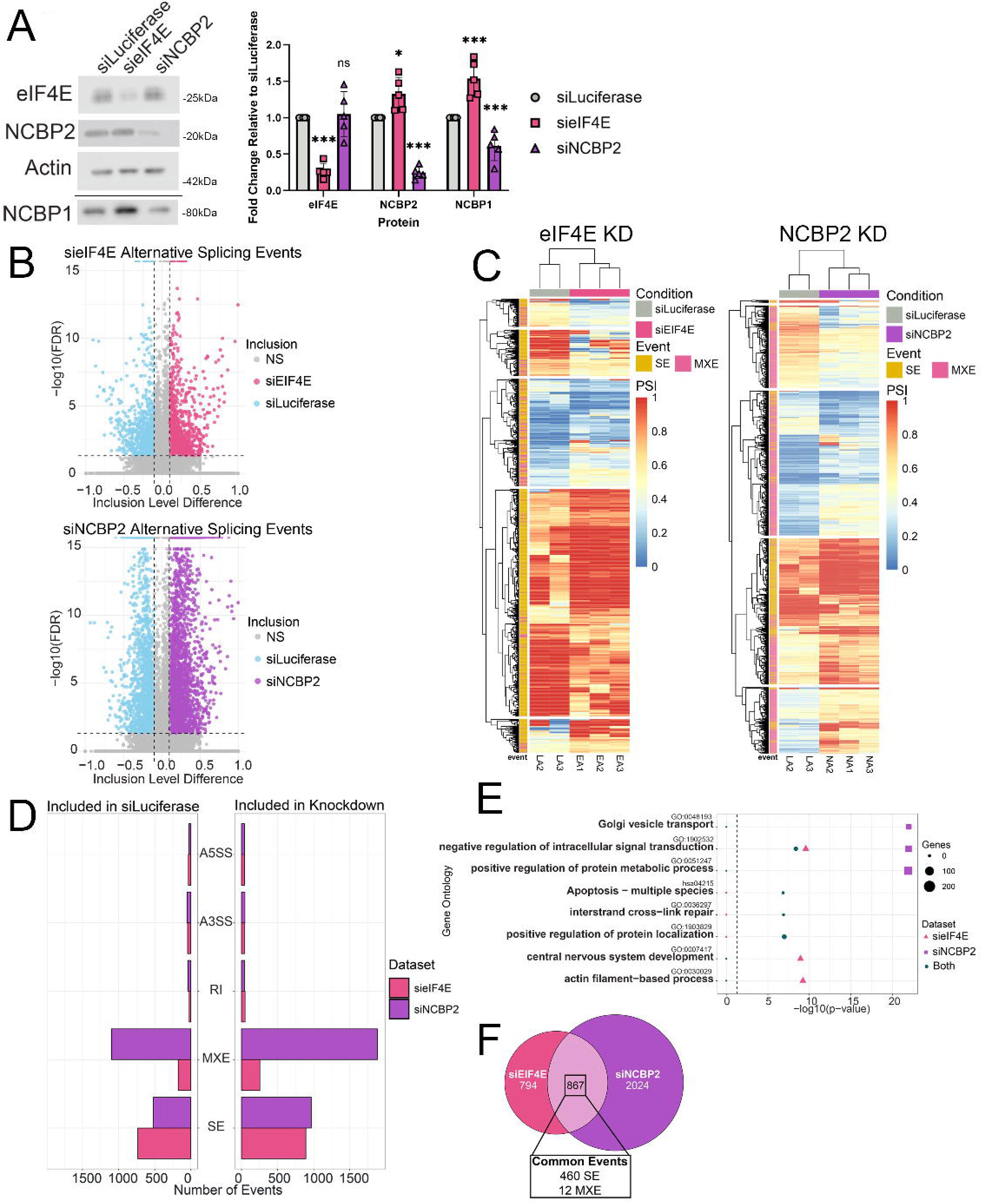
eIF4E and NCBP2 knockdown alter the splicing of distinct targets. **A)** *Left top* Representative western blot of eIF4E, NCBP2, and Actin from U2OS whole cell lysate 96hrs following transfection of siRNA targeting Luciferase, eIF4E, or NCBP2. *Left bottom* Representative western blot of NCBP1 from U2OS whole cell lysate 96hrs following transfection of siRNA targeting Luciferase, eIF4E, or NCBP2. *Right* Quantification of five biological replicates of knockdowns of eIF4E (pink) and NCBP2 (purple) relative to Luciferase (grey). * = p < 0.05, ** = p < 0.01, *** = p < 0.005, ns = non-significant. **B)** *Top* Volcano plot showing the alternative splicing events that were significantly changing in eIF4E knockdown, colored if they are more included in the knockdown (pink) or in control (blue). *Bottom* Volcano plot showing the alternative splicing events that were significantly changing in NCBP2 knockdown, colored if they are more included in the knockdown (purple) or in control (blue). **C)** *Left* Heatmap of significantly changing SE (yellow) and MXE (pink) events showing inclusion (red) or exclusion (blue) of the exon in eIF4E or Luciferase knockdown samples. *Right* Heatmap of significantly changing SE (yellow) and MXE (pink) events showing inclusion (red) or exclusion (blue) of the exon in NCBP2 or Luciferase knockdown samples. **D)** Bar graph illustrating the number of alternative splicing events in EIF4E (pink) and NCBP2 (purple), based if the event is more inclusion in siLuciferase (left) or the knockdown (right). **E)** Dotplot showing the top enriched gene ontology terms in NCBP2 (purple squares), eIF4E (pink triangles), or both (green circles) knockdowns. Size of the point indicates the number of genes and position along the x-axis indicates enrichment of the GO term. Dashed line indicates p = 0.05. **F)** Euler plot comparing the genes that have significantly changing alternative splicing in eIF4E (pink) or NCBP2 (purple) knockdown. The number of SE and MXE events that are changing in both knockdowns are indicated.

We monitored splicing using ribodepletion followed by RNA-Seq as described above (<u>GSE345544</u>). Our sequencing analysis indicated that similar to the case of overexpression, there were no mutations induced by these knockdowns in the splicing factors SF3B1, U2AF1, and SRSF2. Next, we examined the effects of cap-chaperone depletion on the transcriptome via differential expression analysis. Upon genetic reduction of eIF4E, there were 70 genes that were differentially expressed (35 upregulated and 35 downregulated), comprising 1% of the 8306 genes with TPM >10 in the dataset (Figure S2A; Supplemental Table 15). The effects were more striking when NCBP2 was knocked down: 2226 genes were differentially expressed (337 downregulated and 1889 upregulated), comprising 34% of the 6510 genes that had TPM > 10 in the dataset (Figure S2B; Supplemental Table 16). Interestingly, while eIF4E KD has similar magnitude impacts to eIF4E overexpression (6% vs 1%), more transcripts were influenced by NCBP2 KD than NCBP2 overexpression (1.2% vs 34%).

Next, we investigated whether eIF4E and NCBP2 could influence splicing at steady-state. rMATS analysis revealed 2300 alternative splicing events were altered in 1661 genes when eIF4E was knocked down and 4719 alternative splicing events in 2891 genes for NCBP2 using cutoffs of FDR < 0.05 and inclusion level difference <-0.1 or >0.1 (Figure 3B, Supplemental Table 17&18). There is ∼12% overlap between the transcripts alternatively spliced following knockdown of NCBP2 in U2OS and mouse cardiomyocyte cells (Figure S1G), suggestive of a conserved mechanism in regulating alternative splicing. Similar to when cap-chaperones were overexpressed, knockdown led to primarily SE (1627 and 1487 events sensitive to eIF4E and NCBP2 respectively) and MXE (430 in eIF4E and 2982 in NCBP2) events (Figure 3C&D). Similar to overexpression of eIF4E, genetic reduction led to a roughly equal distribution of positive or negative inclusion, while genetic reduction of NCBP2 leads to roughly double the number of SE and MXE events being included than excluded (Figure 3D). In all, 20% of RNAs underwent alternative splicing in eIF4E knockdown, a similar proportion to the overexpression, while in NCBP2 knockdown, a much larger fraction of modified events (44%) underwent alternative splicing compared to overexpression. Thus, eIF4E and NCBP2 influence mRNA processing at baseline as well as overexpression. Indeed, NCBP2, and concomitant loss of NCBP1 (and thus the CBC) loss results in substantially more changes than its overexpression. However, the impacts are still selective, not universal.

Next, we examined whether eIF4E and NCBP2 genetic reduction influenced common or distinct biological pathways. Ontology analysis of alternative splice targets revealed different top GO terms; most enriched in NCBP2 knockdown were “Golgi vesicle transport” and “positive regulation of protein metabolic process” while the top enriched terms in eIF4E knockdown include “central nervous system development” and “actin filament-based process” (Figure 3E, Supplemental Table 19&20). Of the top terms in each dataset only “negative regulation of intracellular signal transduction” was enriched in both knockdown datasets. Therefore, similar to when cap-chaperones are overexpressed, genetic depletion of eIF4E and NCBP2 alters the splicing of transcripts with largely distinct biological functions, in line with the SOCCs model.

Our analysis revealed there were 867 genes that were alternatively spliced in both eIF4E and NCBP2 knockdowns. Of these, 472 alternative splicing events in 408 genes (47% of shared splicing targets) were affected by both knockdowns, 460 of which were SE events and 12 were MXE events (Figure 3F, Supplemental table 21&22). Thus, there are many more shared splicing events upon genetic reduction than overexpression. The genes with these shared events have enriched GO terms “Apoptosis – multiple species”, “interstrand cross-link repair”, “positive regulation of protein localization”, consistent with the known functions of both cap-chaperones^50,51^ (Figure 3E, Supplemental Table 23). The increased number of alternative splicing targets affected by eIF4E and NCBP2 knockdown compared to overexpression indicates that there is a baseline amount of each cap chaperone required for the proper splicing of a class of transcripts, and reduction beyond that threshold alters the splicing dynamics of specific events in those transcripts. Where the baseline is, and what features define the shared events affected by cap-chaperone reduction warrant future investigation.

### Altering cap-chaperone levels is associated with changes in mRNA processing and protein function

To understand the biological consequences of cap-chaperone specification, we examined whether association with a given cap-chaperone was linked to altered mRNA splicing and thus its ultimate functional form. We focused on eIF4E and NCBP2 overexpression relative to vector control cells given their elevation in cancer. Leveraging our above rMATS analysis, we selected a group of targets for further assessment.

Using event-specific primers and RT-qPCR, we observed that *QKI, MAPK8IP3, LRRC23, PLAGL1*, and *ADGRL1* were splicing targets of both eIF4E and NCBP2-dependent splicing as predicted by rMATS (Figure 4A, Table 1). Interestingly for *QKI*, the cap-chaperones had opposing effects: NCBP2 overexpression increased inclusion of exon 3, which encodes the KH domain in QKI protein which binds mRNA; while eIF4E overexpression decreased its inclusion. Another example of modulated mRNA fate comes within the group of mRNAs sensitive to eIF4E but not NCBP2 e.g. *IL1B* and *IL4R*. Using event specific primers, eIF4E overexpression reduced retention of the first intron in *IL1B* and reduced the inclusion of exon 2 in the 5’ UTR in *IL4R* (Figure 4A, Table 1). Within the NCBP2 sensitive group, *VEGFA* and *UBAP2L* were targeted by NCBP2 but not eIF4E, consistent with the rMATS data (Figure 4A, Table 1). *MAPK8IP3* was a target of both eIF4E and NCBP2 overexpression where in both instances elevated cap-chaperone levels led to increased production of transcripts with exon 10 exclusion (transcript variants 1 and 3), i.e. both promoted the same event (Figure 4A, Table 1). We note that none of analyzed transcripts had total mRNA levels altered by either NCBP2 or eIF4E overexpression (Supplemental Figure S2C)^33^.

**Figure 4:**
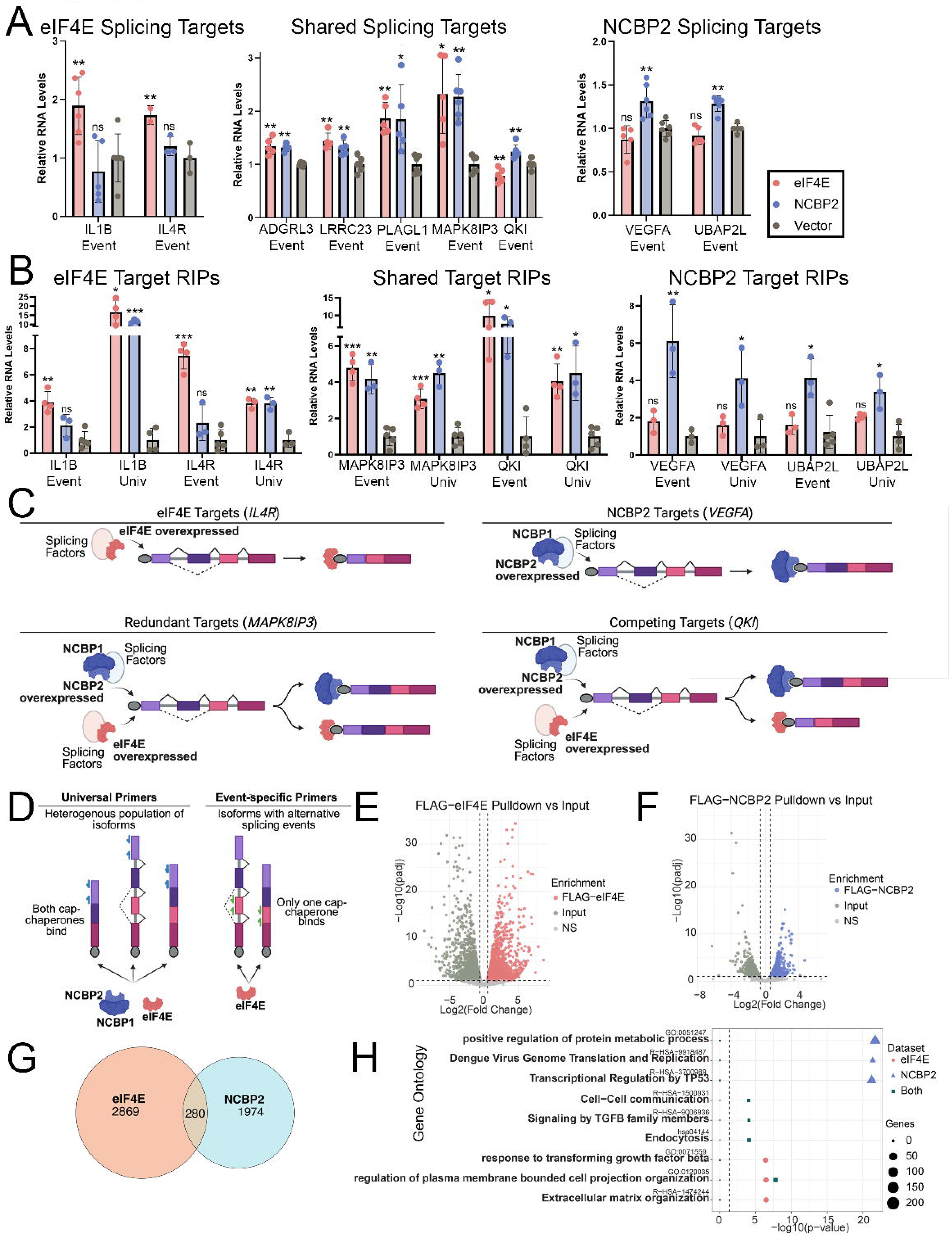
eIF4E and NCBP2 Interact with Distinct RBPs. **A)** Quantitative PCR showing alternative splicing following NCBP2 (blue) or eIF4E (red) overexpression. **B)** Quantitative PCR showing RIPs for alternative splicing targets for FLAG-eIF4E (red) or FLAG-NCBP2 (blue) relative to vector. PCR was conducted with primers specific to the alternative splicing event (event) or common to all isoforms (univ) of a transcript. Note: *IL1B* event-specific primer targets the intron whereas in Figure 4A it targets the spliced exon junction. **C)** Schematic showing the effects of cap-chaperone overexpression on the splicing events of eIF4E targets (top left), NCBP2 targets (top right), redundant targets (bottom left), or competing targets (bottom right) **D)** Schematic of isoforms bound by cap-chaperones when using universal (left, blue) and event-specific primers (right, green). **E)** Volcano plot showing the transcripts that were significantly enriched in the FLAG-eIF4E pulldown (orange) or input (green). **F)** Volcano plot showing the transcripts that were significantly enriched in the FLAG-NCBP2 pulldown (blue) or input (green). **G)** Euler plot showing the overlap between FLAG-eIF4E (orange) and FLAG-NCBP2 (blue) RIP targets from U2OS nuclear lysates. **H)** Dotplot showing the enriched gene ontology terms of transcripts enriched in FLAG-NCBP2 (blue triangles), FLAG-eIF4E (orange circles), or both (green squares) pulldowns. Size of the point indicates the number of genes and position along the x-axis indicates enrichment of the GO term. Dashed line indicates p = 0.05. For all panels, means, standard deviations, and p-values relative to vector (Welch t-test) were calculated in PRISM. * = p < 0.05, ** = p < 0.01, *** = p < 0.005.

Next, we determined whether splicing targets were differentially enriched in cap-chaperones and if this corresponded to the observed splicing outcomes. We conducted RNA immunoprecipitations (RIPs) in nuclear lysates using anti-FLAG antibodies so that differences in antibody avidity would not impact our results and then monitored enrichment relative to nuclear lysates from vector controls by RT-qPCR. The quality of nuclear fractions were verified using Nopp140 and Lamin as nuclear markers and Calreticulin as a cytoplasmic marker (Figure S3A). Importantly, neither eIF4E nor NCBP2 were enriched in the negative control, uncapped 28S rRNA (Figure S3B). For eIF4E-sensitive splicing targets *IL1B* and *IL4R*, FLAG-eIF4E, was enriched in RIPs 3.9 and 7.5-fold respectively using event-specific primers, relative to vector, consistent with our splicing data (Figure 4B, Table 1). By contrast these transcripts were not enriched in FLAG-NCBP2 relative to vector controls in NCBP2 overexpressing cells. In this way eIF4E, but not NCBP2, is positioned to reduce intron retention and thus increase protein production of IL1B, as has been demonstrated following eIF4E activation^52^. Similarly, eIF4E overexpression reduced inclusion of exon 2 in the *IL4R* 5′UTR and increased levels of isoforms 3 and 4, but not isoform 5, likely altering transcript regulation. These specific splice events were enriched in eIF4E but not NCBP2 RIPs; however, using primers found in all isoforms for a given transcript, referred to here as “universal” primers (Figure 4D), we observed that *IL1B* and *IL4R* were found to also immunoprecipitate with NCBP2 (11.8-fold and 3.8-fold respectively) as well as eIF4E (19.2 fold and 3.8 fold respectively) in nuclear lysates (Figure 4B, Table 1). This suggests that a given transcript can be differentially regulated at distinct processing steps by different cap-chaperones (Figure 1A bottom panel & 4D, Table 1). In other words, the cap-chaperone may be switched on the mRNA as part of transcript maturation.

*QKI* presents a particularly interesting case: both eIF4E and NCBP2 RIP the transcript at the same splicing event with similar enrichment relative to vector (7.6-fold for eIF4E, 9.8-fold for NCBP2), and, as expected, both also bound *QKI* using universal primers (4-fold for eIF4E, 4.5-fold for NCBP2) (Figure 4B, Table 1). However, NCBP2 overexpression increased inclusion of the KH RNA-binding domain, while eIF4E overexpression decreased its inclusion, altering the functionality of the protein. This provides an important example whereby competing occupancy by eIF4E and NCBP2 determine splicing outcomes is and protein function (Figure 4C). siRNA knockdown data are consistent with this, where KD of NCBP2 decreased the inclusion of the KH RNA-binding domain similar to eIF4E overexpression (Supplemental Table 17). By contrast, another shared target of eIF4E and NCBP2 RIPs includes *MAPK8IP3* whereby overexpression of either cap-chaperone increases exclusion of exon 10. Using event specific primers, both eIF4E and NCBP2 RIPs were enriched relative to vector (4.2-fold for NCBP2, 4.8-fold for eIF4E; Figure 4B, Table 1), consistent with eIF4E and NCBP2 each being able to influence this splicing event (Figure 4C). Similarly, both eIF4E and NCBP2 were enriched relative to vector using universal primers to *MAPK8IP3* (3-fold and 4.5-fold respectively; Figure 4B, Table 1).

*VEGFA* provides an important example of an NCBP2-sensitive splicing and selective NCBP2 RIP target in nuclear lysates (Figure 4C). NCBP2 overexpression increased exclusion of the heparin-binding domain in *VEGFA*, where this domain enhances the binding of VEGFA protein to the extracellular matrix and increases the more soluble and diffusible form which drive rapid blood vessel growth^53^. At this specific splicing event, NCBP2 bound *VEGFA* with 6.1-fold enrichment relative to vector, compared with 1.8-fold enrichment for eIF4E, the latter not being statistically significant (Figure 4B, Table 1). Using universal primers, *VEGFA* remained primarily an NCBP2 target, with 4.1-fold enrichment for NCBP2 versus a non-significant enrichment of 1.8-fold for eIF4E relative to vector. Thus, *VEGFA* was not enriched in the eIF4E RIP relative to vector and did not induce the same splicing outcome. Similarly, exon 26a *UBAP2L* inclusion is increased by NCBP2 but not eIF4E overexpression, though the functional consequences of increased exon 26a inclusion are not yet known. Here, NCBP2 bound *UBAP2L* with both event-specific and universal primers (4.1-fold and 3.4-fold respectively) while eIF4E did not bind *UBAP2L* more significantly compared to vector using either primer set (1.63-fold and 2-fold respectively) nor did it influence this splicing event (Figure 4A and B, Table 1).

In all, eIF4E and NCBP2 physically interact with their splicing targets as observed using primers specific for relevant splicing events. Using universal primers to assess interactions with the entire population of a given mRNA (Figure 4D), we observe that eIF4E and NCBP2 bind to transcripts that are not affected by the splice event, a prediction of the SOCCs model. In this case, eIF4E or NCBP2 may not mediate the given splice event but influence other nuclear activities such as mRNA export, polyadenylation etc. In all, mRNA fate is influenced in part by which cap-chaperone it is bound during specific processing events, whereby binding of different cap-chaperones to the same transcript are associated with different processing outcomes. These experiments do not rule out additional indirect effects, such as altered splicing factor levels (see below).

### Cap-chaperones associate with distinct but overlapping groups of mRNAs in the nucleus

To establish if eIF4E and NCBP2 physically associate with the same client mRNAs in the nucleus of U2Os cells, we carried out RIP-Seq experiments using anti-FLAG antibodies from nuclear lysates prepared from vector, FLAG-eIF4E, and FLAG-NCBP2 cells (<u>GSE345544</u>). Nuclear quality was assessed using Nopp140 and Lamin as nuclear markers and Calreticulin for the cytoplasm (Figure S3A). Ribodepletion was used for library construction to capture both pre-mRNAs and mRNAs. The use of the FLAG antibody obviated differential binding of eIF4E or NCBP2 antibodies i.e. immunoprecipitation efficiency should be highly similar for FLAG-eIF4E and FLAG-NCBP2. To eliminate off-target binding of the anti-FLAG antibodies, we also analyzed FLAG-vector pulldowns, which only enriched in 12 transcripts relative to input and these were eliminated from the FLAG-eIF4E and FLAG-NCBP2 analysis. We observed no correlation between mRNA abundance and enrichment (Figure S3C). FLAG-eIF4E bound 3149 transcripts when using enrichment cutoffs of fold change > 1.5 and p_adj_ < 0.1 when compared to inputs. This represents 28% of the 11292 transcripts with a TPM > 10 in the FLAG-eIF4E input libraries (Figure 4E, Supplemental Table 24). eIF4E did not RIP with known negative controls mRNAs such as VEGF, GAPDH, and Actin in the nuclear fraction. Consistent with eIF4E’s known functions, gene ontology for the enriched genes include “response to transforming growth factor beta”, “regulation of plasma membrane bounded cell projection organization”, and “extracellular matrix organization” (Figure 4H; Supplemental Table 25). Using the same cutoffs there were 2254 transcripts in FLAG-NCBP2 enriched over input, comprising 22% of the 10267 transcripts with TPM > 10 in the FLAG-NCBP2 input libraries (Figure 4F, Supplemental Table 26). Metascape analysis showed enriched transcripts have gene ontologies including “positive regulation of protein metabolic process” and “transcriptional regulation by TP53” (Figure 4H, Supplemental Table 27). Though both cap-chaperone RIPs are enriched in a similar number of transcripts, they bind largely distinct sets of RNAs, with only 280 transcripts shared between eIF4E and NCBP2 (Figure 4G, Supplemental Table 28). Shared genes have gene ontologies including “signalling by TGFB family members”, “endocytosis”, “response to transforming growth factor beta”, and “regulation of plasma membrane bounded cell projection organization” (Figure 4F, Supplemental Table 29) suggestive of convergent pathways. However, the largely distinct biological processes of genes enriched in FLAG-eIF4E and FLAG-NCBP2 RIP-seq supports the SOCCs model where the cap-chaperones preferentially bind and regulate specific transcripts.

It is important to note that transcripts not enriched in a cap-chaperone RIP does not necessarily mean that the transcript does not interact with that cap-chaperone; while this could be the case for some transcripts, for others it likely indicates that transcripts are less abundant, being under the 10% input levels. Therefore, we measured the quality of our RIP-seq libraries in order to determine how abundant the enriched targets were. We used the following metric: median transcripts per million minus the median absolute deviation (median [TPM]–MAD). In the eIF4E RIP-seq we observed 8703 transcripts with score > 10 and 632 with score > 100, while ∼70% of the transcripts had a score < 1. Similarly, in the NCBP2 RIP-seq we observed 9003 transcripts with score > 10 and 505 with score > 100, while ∼69% of the transcripts had a score < 1. Thus, while this experiment allowed us to identify binding targets of eIF4E and NCBP2 in U2OS cells with similar efficiency, the overall sensitivity was limited and could underrepresent the number of eIF4E- and NCBP2-bound nuclear transcripts. Despite these limitations, our data showed that NCBP2 and eIF4E preferentially interact with distinct but overlapping sets of client RNAs.

### eIF4E physically associates with sites of active transcription

We show that in the nucleus eIF4E and NCBP2 impact different but overlapping splicing events and similarly bind to distinct but overlapping mRNA populations. These differences could arise due to spatial segregation of cap-binding chaperones associated with co-transcriptional and post-transcriptional splicing. Indeed, NCBP2 is well known to physically associate with sites of active transcription^6^ while the role for eIF4E there was unknown. With this in mind, we examined whether endogenous eIF4E associated with transcription sites using antibodies directed against RNA polymerase II phosphorylated at Serine 2 (POLR2A-S2P) to mark active transcription^54,55^. eIF4E immunoprecipitations from nuclear lysates revealed interactions between endogenous eIF4E and POL2RA-S2P (Figure 5A). eIF4E interacted with positive controls HuR/ELAV1, SF3B1 and U2AF1 but not the negative control Lamin; nor were any of these proteins found in the IgG controls. H2B and Lamin served as fractionation controls for the nucleus and MEK1 and HSP90 for the cytoplasm (Figure 5A&B).

**Figure 5.**
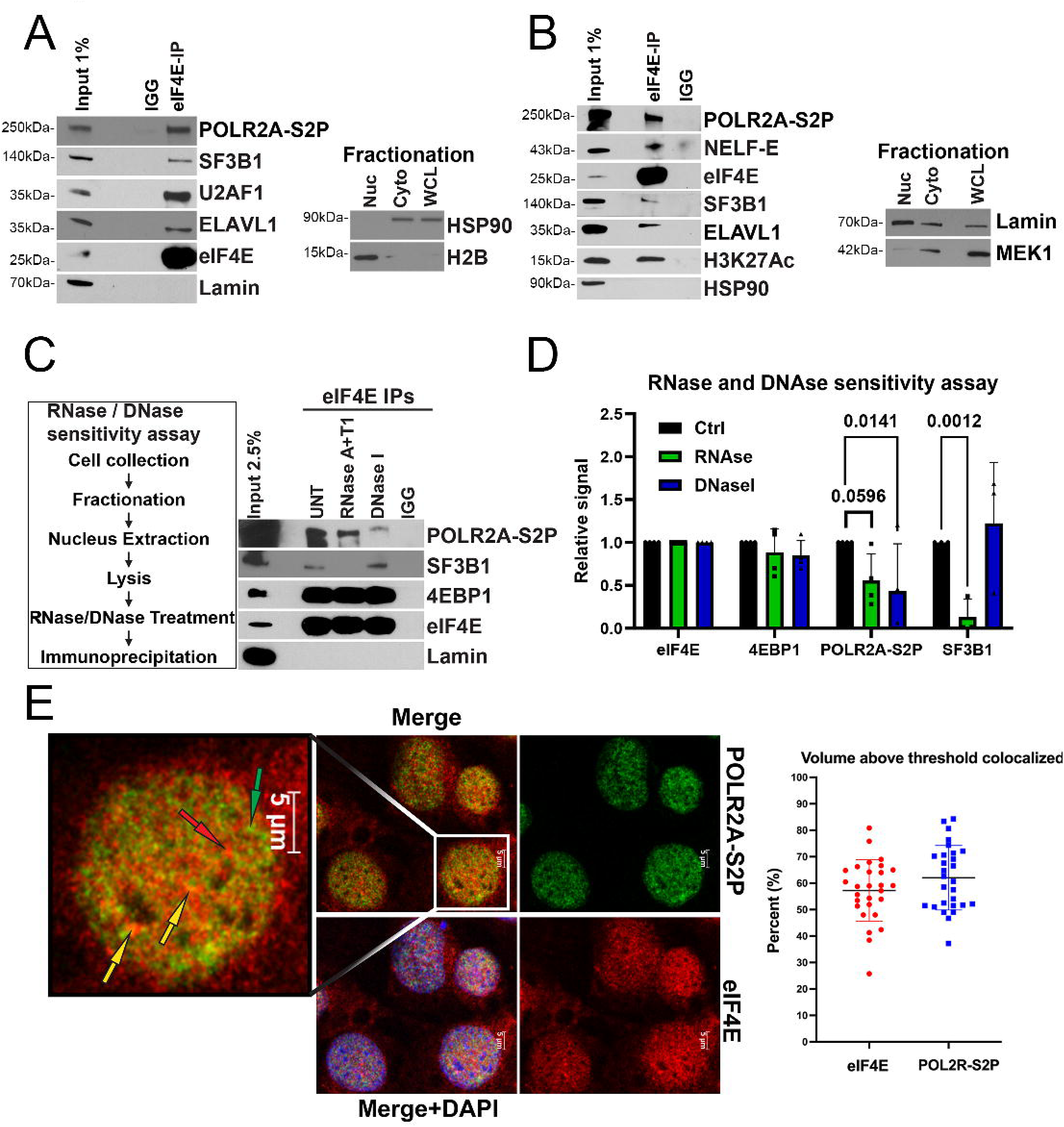
A population of eIF4E interacts with active sites of transcription. **A)** Endogenous eIF4E co-immunoprecipitated with POLR2A-S2P in the nuclear fraction of U2OS cells. Immunoprecipitations (IP) were carried out using nuclear lysates and anti-eIF4E antibody (eIF4E-IP) or appropriate IgG control (IGG), carried out at least three independent times with one representative experiment shown. Samples were analyzed by western blot (WB) using antibodies as indicated. Fractionation controls for HSP90 for the cytoplasmic fraction (Cyto), histone 2B (H2B) for the nuclear fraction (Nuc) are shown. Whole Cell lysis (WCL). **B)** Endogenous eIF4E co-immunoprecipitated with POLR2A-S2P in crosslinked nuclear fractions using NEXSON, one representative experiment is shown of three biological replicates. **C)** The impact of DNAse and RNAse on the eIF4E-POLR2A-S2P interaction were analyzed by western blot. **D)** Quantification of three or more biological replicates from C) showing the relative signal of each protein in control (black), RNase-treated (green), and DNase-treated conditions (blue) (p-values are provided using ANOVA in PRISM, all comparisons not indicated were non-significant). **E)** *Left* Confocal micrographs of U2OS cells stained with antibodies against eIF4E (red) and POLR2A-S2P (green), with nuclei counterstained with DAPI (blue) and overlay in yellow (merge). Single section through the plane of the cell is shown. Arrows indicate examples of eIF4E-only (red), POLR2A-S2P-only (green), or co-localized (yellow) signals. *Right* Quantitation of colocalization using Imaris, data were collected as Z-stacks for >25 cells. Percentage colocalization over the cell volume was calculated in Imaris and processed in PRISM. Each data point represents a single nuclei.

To ensure that the eIF4E-POL2RA-S2P interaction did not arise due to reassortment during immunoprecipitation, we employed the nuclear extraction by sonication method (NEXSON)^56^ which involves crosslinking cells prior to isolation of nuclei by sonication and additionally better solubilizes the transcription machinery (Figure 5B). Like the above native (not crosslinked) nuclear lysates, we observed that endogenous eIF4E bound to a fraction of POLR2A-S2P but not to the negative control. We also observed that eIF4E bound to another marker of active transcription, acetylated histone H3K27 (H3K27Ac)^57^ and NELF-E^58,59^, a component of the RNA polymerase complex responsible for pausing, which is important for coordinating transcription with splicing^5,6,59^ (Figure 5B).

To assess whether eIF4E required intact chromatin and transcripts for this physical interaction, we monitored DNAse and RNAse sensitivity of the eIF4E-POL2RA-S2P interaction in uncrosslinked nuclear fractions. Nuclear lysates were treated with DNAse I or RNAses T and A1 followed by eIF4E immunoprecipitation. Representative blots and quantitation are shown (Figure 5C&D). DNAse I treatment reduced the eIF4E-POL2RA-S2P interaction by ∼11-fold relative to untreated controls and RNAse treatment by ∼1.5 fold. eIF4E directly interacts with 4EBP1 and this interaction was not disrupted by either DNAse or RNAse, as expected. These factors were not observed in the IgG controls and Lamin served as a negative control. Immunoprecipitation efficiency between experiments was controlled by normalizing to the efficiency of eIF4E immunoprecipitation of itself in the same nuclear lysate, which we note were not impacted significantly by either DNAse or RNAse (Figure 5D). Finally, the interaction of eIF4E with SF3B1 was RNAse-dependent as shown previously^33^ and interestingly, not reliant on DNA as seen by insensitivity of the eIF4E-SF3B1 interaction to DNAse I treatment (Figure 5C&D). In all, the physical interactions between eIF4E and POL2RA-S2P depended on the presence of DNA and RNA consistent with a reliance on intact transcription complexes and proximity to mRNA processing events. We note that similar to eIF4E, interactions between NCBP2 and splicing factors, including U2AF1, SNRNP200, PRPF4, and SNRNP70 are RNA-dependent^38^. Similar to the RNase sensitivity of the eIF4E and POL2RA-S2P interaction, the association of the CBC, measured by monitoring NCBP1, and RNA polymerase II during splicing was shown to be RNA dependent^60^. Thus, eIF4E and NCBP2 similarly engage the splicing and transcription machinery in an RNA-dependent manner.

To ensure the interaction could be observed in intact cells, we employed immunofluorescence and confocal laser microscopy collecting Z-stacks throughout the cell volume. We observed that eIF4E is localized in spheroid nuclear structures as well as diffusely throughout the nucleoplasm and cytoplasm consistent with previous studies ^13,15,17,18,61^ (Figure 5E). POL2RA-S2P was similarly found in puncta as well as a diffusely throughout the nucleus and excluded from the cytoplasm. Quantification of colocalization within the nucleus was performed using Imaris. We observed that 57 ± 11% of eIF4E co-localized POL2R-S2P and conversely 62 ± 12% of POL2RA-S2P colocalized with eIF4E (Figure 5E). The average original Mander’s overlap coefficient which provides the correlation of absolute intensity was 0.74 for eIF4E and >0.9 for POL2RA-S2P in eIF4E/POL2RA-S2P co-staining, indicating this was non-random (Figure S3D). Note that for quantitation, secondary antibody staining served as the background signal for subtraction. In all, we observed that a fraction of endogenous eIF4E physically associated with transcriptionally active RNA polymerase II and other markers of active transcription.

### Physical separation of eIF4E and NCBP2 may contribute to cap-chaperone specification

Given that a fraction of both eIF4E and NCBP2 were found at active sites of transcription, we explored whether differential influence on mRNA fates could arise from partial separation of eIF4E and NCBP2 in the nucleus at different transcription sites. To assess spatial association, we monitored localization of eIF4E and NCBP2 by confocal microscopy (Figure 6A). We observed that 46 ± 18% of NCBP2 with eIF4E and 10 ± 8% of eIF4E with NCBP2 (Figure S3E). Thus, eIF4E and NCBP2 existed mostly separately but could be found together. Indeed, endogenous eIF4E immunoprecipitated with NCBP2 and its co-factor NCBP1 in nuclear fractions from U2OS cells (Figure 6B&C).

**Figure 6.**
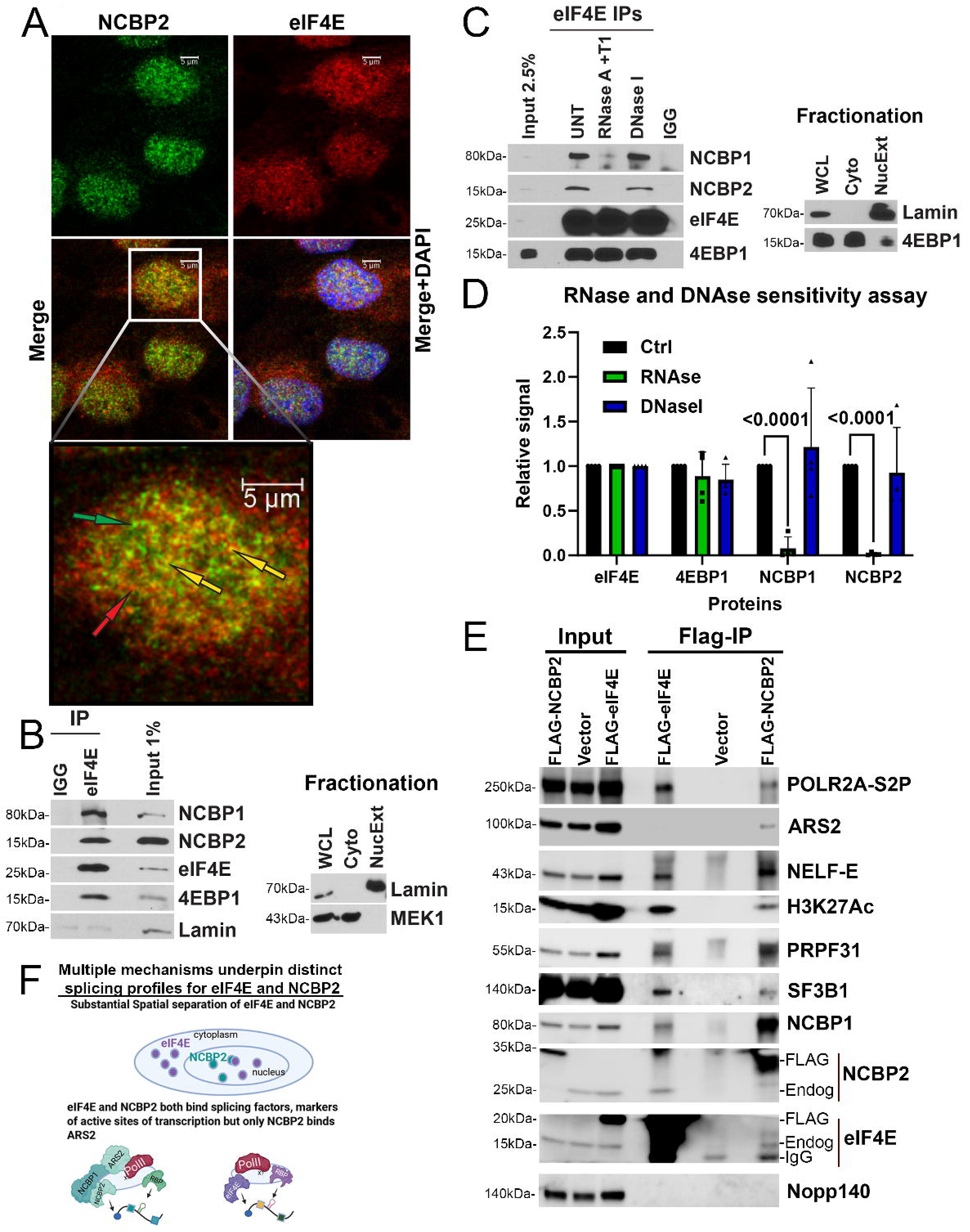
Mechanisms for eIF4E and NCBP2 dependent splicing. **A)** Confocal micrographs of a single section through the plane of U2OS cells stained with antibodies against endogenous eIF4E (red) and NCBP2 (green), overlay (yellow) and nuclei with DAPI (blue). Arrows indicate examples of eIF4E-only (red), NCBP2-only (green), or co-localized (yellow) signals. **B**) Endogenous eIF4E co-immunoprecipitated with NCBP1 and NCBP2 in nuclear fractions from U2OS cells. Immunoprecipitations (IP) were carried out using U2OS nuclear lysates and anti-eIF4E antibody (eIF4E-IP) or appropriate IgG control (IGG), carried out at least three independent times, with a representative experiment shown. Fractionation controls included Lamin (nuclear fraction, Nuc), MEK1 served as controls for the cytoplasmic fractions (Cyto). Whole Cell lysis (WCL). **C)** The impact of the RNAse and DNAse treatment on of the interaction between endogenous eIF4E and NCBP2 and NCBP1. IPs were performed using nuclear lysates from U2OS cells with or without RNase A/T1 or DNase I treatment prior to IP (as in Figure 5C). IP samples, along with input, were analyzed by western blot using the indicated antibodies. **D)** Quantification of three biological replicate experiments showing the relative signal of each protein in control (black), RNase-treated (green), and DNase-treated conditions (blue) (p-values are provided using ANOVA in PRISM, all comparisons non indicated were non-significant). **E)** FLAG IPs for nuclear lysates of NCBP2- or eIF4E-overexpressing cells showing overlapping but distinct complexes were formed. Representative experiments of two different eIF4E and NCBP2 clones in biological replicates are shown. Vector control cells and probing for NOPP140 served as negative controls. **F)** Model of the mechanisms that underpin the distinct splicing profiles of eIF4E and NCBP2. These include spatial separation of the proteins (top) and interaction with different binding parters (bottom).

No immunoprecipitation was observed for Lamin, and no interactions were observed in the IgG controls. NCBP1 and NCBP2 interactions with eIF4E were reduced by 27-fold and 10-fold upon treatment with RNAse, respectively (Figure 6C&D). By contrast, DNase I treatment did not significantly reduce the binding of eIF4E to either NCBP1 or NCBP2. The interaction between eIF4E and 4EBP1 was insensitive to either DNAse or RNAse as expected given its direct interaction with eIF4E^33^ providing a negative control. Thus, the eIF4E-NCBP2/NCBP1 interaction was RNA-dependent suggestive that these cap-chaperones localized to sites of mRNA processing, such as transcription or splicing factories, which are disrupted by RNAse treatment but did not require DNA. This finding also indicates that NCBP1 is not directly binding eIF4E to modulate cap-binding activity as it does for NCBP2, since direct binding would not be abrogated by RNAse treatment. In all, the largely distinct localization of eIF4E and NCBP2 likely contribute to their distinct modulation of transcriptomic and splicing signatures.

### eIF4E and NCBP2 bind common transcriptional machinery but only NCBP2 binds the ARS2 adaptor

Several key factors are known to underpin the capacity of the NCBP2/NCBP1 complex to associate with the processing machinery including ARS2, also known as SRRT^6^.

ARS2 mediates interactions between NCBP2/NCBP1-capped mRNA complexes early in transcription, playing a role in recruiting many factors including the splicing and mRNA export machinery to the NCBP2/1 complexes bound to mRNAs^6,24^. We examined the possibility that ARS2 was specifically interacting with NCBP2 and not eIF4E complexes as another possible mechanism underlying their distinct impacts on splicing.

Interactions were examined in two different eIF4E and NCBP2 stable cells lines each in biological duplicate and crosslinked nuclei to avoid reassortment (Figure 6E).

Calreticulin and Cytochrome C served as the cytoplasmic and Lamin as the nuclear fractionation control (Figure S3F). We observed that FLAG-eIF4E interacted with NCBP1 and NCBP2 as well as NELF-E and H3K27Ac as seen with the endogenous splicing machinery (Figures 5&6). As expected, FLAG-NCBP2 was complexed to ARS2, NELF-E, eIF4E and POL2RA-S2P^6^. Interestingly, FLAG-eIF4E did not interact with ARS2. Thus, the ARS2 containing mechanism utilized by NCBP2 to interact with the splicing machinery is not shared between the cap-chaperones. In this way, these factors have distinct partner protein configurations that may underlie the different splicing landscapes.

### NCBP2 and eIF4E differentially influence production of the splicing machinery

Previously it was shown that eIF4E manifested splicing changes through two simultaneous effects: its physical association with some splicing substrate mRNAs and the splicing machinery as we showed above, as well as enhanced production of some components of the splicing machinery including SF3B1, U2AF1, SNRNP200 and PRPF31^33^. Here, we examined if NCBP2 similarly impacted splice factor production. For this comparison it is important to note that eIF4E and NCBP2 overexpression relative to respective vector controls were both ∼3.5-fold and thus comparable (Figure 1C).

Notably, this level of overexpression is in the mid-range of expression seen in eIF4E and NCBP2 overexpression cancers and is thus physiologically relevant^40,62^. eIF4E increased protein production of its mRNA export and translation targets: MYC by ∼2.5-fold and MCL1 by ∼2.2-fold relative to vector while NCBP2 did not alter production of these (Figure S4A). Interestingly, NCBP2 did not increase production of U2AF1, SNRNP200 or PRP31 relative to vector controls while eIF4E elevated these by ∼2-fold consistent with previous work^33^ (Figure S4A). NCBP2 elicited an increase in SF3B1 protein levels by ∼1.7-fold relative to vector while eIF4E has ∼2-fold change consistent with previous studies^33^. Also, eIF4E overexpression did not alter the levels of UsnRNAs by RT-qPCR consistent with previous studies^33^, though NCBP2-overexpression modestly increased U5 snRNA ∼1.5-fold compared to vector (Figure S3E). In all, these two cap-chaperones produced different splicing factor landscapes which likely contributed to their distinct splicing signatures. Thus, in addition to their modulation of splicing through association with client RNAs described above, eIF4E and NCBP2 are positioned to also influence splicing indirectly.

## Discussion

Direction of nuclear mRNA fate through the m^7^G cap was considered the sole purview of NCBP2 (Figure 1A, top panel). This model was based upon the assumption that NCBP2 was the only nuclear cap-binding protein and that its activity was a simple housekeeping function (Figure 1A, top panel). However, endogenous eIF4E and NCBP2 co-exist in the nuclei of plant, yeast and animal cells^9,10,13–15,39^ and both bind the m^7^G cap in nuclear lysates and physically associate with 1000s of mRNAs consistent with previous studies^17,29,49^ (Figure 1B, Supplemental Figure 1A, Suplemental Tables 24 & 26). Interestingly, we observed that eIF4E and NCBP2 overexpression or knockdown resulted in altered splicing for 1000s of transcripts (Figure 2C & 3F). These transcripts were largely distinct with only a small percentage of overlapping splice events modulated by both: with 1-3% alternative splicing events in common in eIF4E and NCBP2 overexpression and 3-25% events in common upon genetic reduction. Thus, the predominant purpose for coexistence of cap-chaperones in the nucleus is unlikely to be functional redundancy.

Therefore, we propose that cap-chaperones can select specific mRNAs for processing, thereby embodying a novel regulatory control point as outlined in the SOCCs model (Figure 1A, bottom panel). In support of this possibility, we observed that changing cap-chaperone levels is accompanied by altered cap-chaperone association and processing outcome, with consequences for function (Figure 4, Table 1). For example, using event specific primers, we showed that *QKI* inclusion of exon 3 which encodes its KH RNA binding domain is dependent on relative levels of eIF4E and NCBP2. NCBP2 overexpression led to increased inclusion of exon 3, using event-specific primers NCBP2 overexpression led to a 9.8-fold enrichment of *QKI* exon 3 containing pre-mRNAs relative to vector controls. By contrast while eIF4E overexpression increased association of the same *QKI* exon 3 containing pre-mRNA with the eIF4E protein relative to vector, eIF4E overexpression reduced inclusion of exon 3 which in turn will increase production of QKI protein without its RNA binding domain. Importantly, *QKI* RNA levels did not change upon eIF4E or NCBP2 overexpression^33^ (Supplemental Figure 2C). QKI acts as a tumor suppressor in glioblastoma, non-small lung cancer, prostate, pancreatic, colorectal, bladder, breast cancer and hepatocellular carcinoma which all showed decreased expression of QKI^63^. Loss of function of the KH domain alters QKI’s functions in alternative splicing, mRNA stabilization, and transport^64^ and mutation of the domains is embryonic lethal in mice^65,66^.

We also observed instances where only one cap-chaperone RIP was enriched in a given pre-mRNA and this elicited altered mRNA splicing and thus will alter biological outcomes. For instance, eIF4E overexpression increased association of pre-mRNAs containing the first intron of *IL1B* with eIF4E leading to reduced intron retention in these transcripts; conversely NCBP2 overexpression did not increase NCBP2’s association with *IL1B* nor did it modulate levels of intron retention relative to vector (Figure 4A&B). Indeed, eIF4E mediated reduction in intron retention is consistent with studies demonstrating that eIF4E overexpression leads to increased IL1B protein production^52^.

IL1B promotes survival and motility in U2Os cells, consistent with it being a downstream effector of eIF4E^67,68^. *VEGFA* splicing is sensitive to NCBP2 and not eIF4E overexpression and is enriched in NCBP2 but not eIF4E RIPs relative to vector controls. NCBP2 increased exon 6 exclusion of *VEGFA* which encodes the heparin-binding domain. Functionally speaking, the exclusion of the exon 6 coding heparin-binding domain significantly decreases binding with the heparin sulfate proteoglycan in the extracellular matrix, making the growth factor mostly soluble and allowing formation of the gradient that will promote formation of capillary branched networks and angiogenesis to support tumor growth^69^. *MAPK8IP3* provides a case of redundancy, where exon 10 exclusion is increased and exon 10 containing pre-mRNA becomes enriched in either eIF4E or NCBP2 upon their respective overexpression. Through interaction with the JNK signaling pathway, MAPK8IP3 has been linked to cellular response to stress and cellular transport^70^, which is consistent for being a downstream regulator of the oncogenic properties associated with overexpression of eIF4E and NCBP2. In all, our data indicate that eIF4E and NCBP2 are associated with differential effects on mRNA splicing events that are positioned to impact protein structure and cell physiology, supporting cap-chaperone specification as a candidate regulatory node.

Our above studies indicate that cap-chaperones associated with a particular pre-mRNA isoform can be altered i.e. that mRNAs may undergo cap-chaperone switching. Given this, we examined whether the population of particular transcripts could be associated with either cap-chaperone using universal primers i.e. primers that capture all the pre-mRNA and mRNA isoforms for that transcript (Figure 4D). Importantly, each capped transcript can only be bound by either eIF4E or NCBP2 at any given time. Using universal primers and nuclear lysates, we observed in some transcript populations that either eIF4E or NCBP2 overexpression led to increased enrichment for the population in respective eIF4E or NCBP2 RIPs (Figure 4B). Importantly, within these transcript populations, mRNAs are undergoing various processing steps e.g. splicing, polyadenylation, export etc. Indeed, eIF4E and NCBP2 interact with these machineries^17,21,26,27,29,31–33,71–74^. Thus, eIF4E and NCBP2 sample transcript populations across processing steps. In this way, binding of eIF4E and NCBP2 to mRNAs that are not splicing targets is expected and may reflect cap-chaperone switching between processing steps as well as among specific events. For instance, eIF4E immunoprecipitated *CCND1* and *MYC* mRNAs to stimulate their nuclear mRNA export but did not influence their splicing. Another example involves the enrichment of first-intron containing *IL1B* by eIF4E but not NCBP2 (Figure 4B, Table 1). However, overexpression leads to increased binding of either NCBP2 or eIF4E to *IL1B* using universal primers (Figure 4B, Table 1). In this case, the binding of NCBP2 is not related to splicing of the first intron of *IL1B* given its absence from RIPs with event-specific primers and lack of impact on this splicing event (Figure 4A&B, Table 1). Thus, NCBP2 is likely involved in some other processing step e.g. polyadenylation, mRNA export etc. Another case demonstrates preferable binding of NCBP2 upon NCBP2 overexpression for both event specific events and universal primers for *VEGFA* and *UBAP2L* while eIF4E is not enriched for binding using either sets of primers upon eIF4E overexpression. In this latter case, it appears eIF4E is not involved substantially in the processing of these transcripts. In all, at different stages of processing mRNAs can undergo cap-chaperone switching for regulation of specific splicing events or processing/export steps.

Our dissection into the molecular basis of mRNA selectivity and subsequent generation of distinct splicing profiles suggests that there may be multiple, contemporaneous mechanisms at play (Figure 6F). One possibility was that eIF4E could be limited to post-transcriptional splicing and NCBP2 to co-transcriptional splicing. Surprisingly, we observed that eIF4E bound to transcriptionally active RNA Pol II and other transcription factors; thus, eIF4E and NCBP2 are both found at active transcription sites (Figure 5).

However, we did note that there was a substantial spatial separation of eIF4E and NCBP2 by confocal microscopy which likely underpins at least in part the different splicing profiles and transcript preferences (Figure 6A). We also observed differences in protein complexes formed with each chaperone in the nucleus, where ARS2 only associated with NCBP2 (Figure 6E), suggesting that each cap-chaperone could have distinct cohort of protein interactors to select mRNA clients and recruit to specific mRNA processing machineries. Our studies also provided evidence for eIF4E and NCBP2 to differentially modulate splice factor production which could influence splicing even for mRNAs not physically associated with a given cap-chaperone (Figure S4). Finally, eIF4E overexpression did not substantially alter NCBP1 or NCBP2 protein levels (Figure 1C), whereas NCBP2 knockdown reduced NCBP1 and eIF4E knockdown modestly increased NCBP1 and NCBP2 (Figure 3A); cross-regulation may also occur in the knockdown experiments. Importantly, this selectivity does not appear to be based on relative affinities since eIF4E and NCBP1/NCBP2 use similar structural strategies to bind the m^7^G cap with similar affinities in vitro^21,23,24^ and similar enrichments on cap-columns using nuclear lysates (Figure 1B). In all, we identified multiple features that may in aggregate contribute to differential specification of mRNAs to a given cap-chaperone (Figure 6F).

The traditional view predicts that modulation of NCBP2 would influence splicing globally (Figure 1A top panel); however, we observed only 15-30% of splice events were modulated by NCBP2 overexpression. Consistent with these results, recent reports demonstrated that NCBP2 overexpression influenced the splicing of only ∼1000 transcripts in mouse cardiac cells^37^, 155 of which were in common with those found here (Figure S1F). Similarly, knockdown of NCBP2 in mouse cardiomyocytes leads to alternatively splicing of ∼1000 transcripts^37^, 365 were also alternatively spliced in our NCBP2 knockdown (Figure S1G). Similar to our findings, this study showed that NCBP2 principally impacted SE events which only rarely corresponded to altered transcript levels^37^. The splicing effects of NCBP2 in cardiac cells were sufficient to retard heart maturation *in vivo* demonstrating the biological relevance of this pathway^37^. While eIF4E elevation is well known to contribute to multiple cancer phenotypes^25,75,76^, recent studies have also implicated increased NCBP2 levels as a contributor to malignancy^62,77,78^ supporting the biomedical relevance of this regulatory point. This also suggests the possibility that more nuclear cap-chaperones exist and can regulate the fate of specific mRNAs. Indeed, the recent discovery of NCBP3 as another nuclear cap-binding protein^5^ is easily incorporated into the SOCCs model and suggests that understanding the chaperoning of capped-RNAs through the gene expression steps may be a multi-factor rather than a two-factor problem. Indeed, our data suggest mRNAs may even be exchanged between cap-chaperones at different stages of processing consistent with our observations that transcripts can be detected on a given cap-chaperone with universal primers but not always with splice event specific primers (Figure 4B, Table 1).

Together, our findings support SOCCs as a candidate layer of nuclear mRNA regulation in which eIF4E and NCBP2 engage distinct, but overlapping, mRNA populations (Figure 1A). Rather than acting as interchangeable carriers of the m^7^G cap, these factors may help determine the molecular context in which transcripts are processed. In all, cap chaperones could be a key component to establish the specific molecular environment around each mRNA molecule. Driven by their localization, their affinity for cap, their interaction with other nuclear agents (splicing factors, polyadenylation factors, transcription factors), the regulation of local environment (open chromatin, active transcription), the abundance of their targets and their own relative abundance (competition), the role of cap-chaperones could be to create a very adaptable environment for mRNA maturation. This view also raises the possibility that SOCCs contributes to the spatial organization of mRNA processing within dynamic nuclear assemblies. Incorporating this model into our current understanding of gene expression will better enable us to target cap-chaperone proteins and predict the impacts of targeting these in patients both in terms of potential compensatory mechanisms and biological impacts. In all, our findings support directed cap-chaperoning of transcripts which incorporates a cap-chaperone specification step to influence mRNA fate with biological impacts.

## Materials and Methods

### Cell culture

U2OS cells were obtained from ATCC and maintained at 37°C and 5% CO2 in Dulbecco’s modified Eagle’s medium (DMEM) (ThermoFisher Scientific) supplemented with 10% fetal bovine serum (FBS) (ThermoFisher Scientific) and 1% penicillin– streptomycin (ThermoFisher Scientific). pcDNA-2FLAG-eIF4E and vector stable cell lines were generated previously^16,29,27^, and maintained as U2OS cells with the addition of G-418 (1 mg/ml, Wisent Bioproducts). NCBP2-pRETS was a kind gift of A Segref and I Mattaj. pcDNA-2FLAG-NCBP2 was generated and included the entire coding region of NCBP2 confirmed by DNA sequencing. Stable FLAG-NCBP2 cell lines were generated as described^16,29,27^. Cell lines were authenticated using STR profiling (Wyndham Forensic Group). Cultured cells were routinely checked to ensure that there was no mycoplasma contamination by PCR^79^. Knockdowns were conducted in vector stable cells using 50nM of siRNA (siLuciferase and sieIF4E sequences as described previously^80^; siNCBP2 is Hs_NCBP2_8 5’-CAGGACTACGATGCTGGGAGA-3’ from Quiagen) using lipofectamine RNAiMax (ThermoFisher) for 96 hours.

### Western blot analysis

Western blot analysis was performed as described previously^80^. Blots were blocked in 5% milk in TBS–Tween 20. Primary antibodies were diluted in 5% milk. Quantification of blots was carried out using FIJI and plotted in PRISM. Antibodies used are found in Supplemental Table 30.

### RNA extraction and quantitative PCR

DNAse treated RNA samples (Direct-zol RNA Miniprep Kit, Zymo Research) were reversed transcribed using MMLV reverse transcription (ThermoFisher Scientific), qPCR analyses were performed using SensiFastSybr Lo-Rox Mix (Bioline) in Applied Biosystems QuantStudio thermal cycler using the relative standard curve method (Applied Biosystems User Bulletin #2). All conditions were previously described^33^. Primers used are available in Supplemental Table 31.

### RNA Sequencing and Differential Gene expression analysis for total RNAs

RNAs isolated from nuclear extracts from six FLAG-NCBP2 and three FLAG vector stable cell lines underwent ribodepletion using the KAPA RNA HyperPrep Ribodepletion Kits (Roche) according to the manufacturer’s instructions. Overexpression libraries were subjected to flow cell high-output (75 cycles) singe-end sequencing on an Illumina NextSeq 2000 Sequencer at the genomics facility at the Institute for Research in Immunology and Cancer (IRIC) at the University of Montreal. Overexpression RNA-seq data are deposited as <u>GSE300496</u>. We note that CBC1A1 of the overexpression libraries was removed from the analysis due to issues with library preparation that introduced mutations, leaving five NCBP2 overexpression and three vector datasets.

Knockdown and RIP-seq libraries were sequenced by Admera Health on an Illumina NovaSeq X Plus sequencer. For both eIF4E and NCBP2 we analyzed three knockdown samples and compared to two siLuciferase samples. Knockdown RNA-seq data are deposited as <u>GSE345544</u>.

RNA-sequencing reads were trimmed using Trimmomatic version 0.35^81^ and aligned to the human genome GRCh38 using STAR version 2.7.1a^82^ with gene annotations from Gencode version 32 to match previous data for eIF4E. Differential expression analysis was conducted using DESeq2 version 1.30.1^83^. Genes were considered to be significantly differentially expressed if they were changing more than 2-fold (log_2_FoldChange > 1) and had an FDR-adjusted p-value (p_adj_) < 0.01 unless otherwise noted.

### Alternative Splicing Analysis

Alternate splicing analysis was conducted on the STAR-aligned datasets using rMATS turbo version 4.3.0^84^ using the following settings: -t paired –nthread 20 –readLength 101 –variable-read-length –libType fr-unstranded. Gencode version 32 transcripts with transcript support level 1 were used for annotation of splice events. For the following analyses using the eIF4E U2OS rMATS analysis^33^, alternate splicing events were considered significant if FDR < 0.15 and the absolute inclusion level difference > 0.05; while for the new NCBP2 overexpression and NCBP2 and eIF4E knockdown datasets events were considered significant if FDR < 0.05 and the absolute inclusion level difference > 0.01. The inclusion level difference of eIF4E overexpression rMATS was multiplied by -1 to make it consistent with the other rMATS data, where a positive inclusion level difference indicates higher inclusion in the overexpression or knockdown samples. Coordinates of the skipped exons undergoing significant changes in each group were extracted from the rMATS results. Introns surrounding the skipped exons were defined by the sequence between the skipped exon and the upstream exon end coordinate or downstream exon start coordinate as recorded in the rMATS output.

Shared SE events were matched by identifying events with identical gene names, exon start sites and exon ends while shared MXE events were identified based on gene name and start coordinates of the two mutually exclusive exons.

### Position analyses of events in transcripts

Previously, we normalized location by the total number of exons so that the x-axis represented the location in percent of the total gene for an exon. Here, we report the x-axis by the actual exon number. For SE events, the location of the skipped exon is reported. The distribution of skipped exon position was compared to all exons in the GTF used for rMATS analysis using a χ^2^-goodness of fit test.

### Cellular fractionation

About 5 × 10^7^ U2OS cells were collected and washed twice in ice-cold PBS (300 × g for 3–5 min) and then resuspended with slow pipetting in 0.5 ml of lysis buffer B (10 mM Tris (pH 8.4), 140 mM NaCl, 1.5 mM MgCl2, 0.25% Nonidet P-40, 1 mM DTT, 100 U/ml RNase inhibitors). The lysate was centrifuged at 1,000 × g for 3 min at 4°C, and the supernatant (cytoplasmic fraction) was transferred into a fresh microtube. The pellet (nuclear fraction) was resuspended in 1 Volume of lysis buffer B and transferred to a round-bottomed polypropylene tube, and 1/10 volume of detergent stock (3.3% sodium deoxycholate, 6.6% Tween 40 in DEPC H2O) was added with slow vortexing (to prevent the nuclei from clumping) and incubated on ice for 5 min. The suspension was transferred to a microtube and centrifuged at 1,000 × g for 3 min at 4°C. Supernatant (post-nuclear fraction) was added to the cytoplasmic fraction.

### Immunoprecipitation with and without RNAse and DNAse treatment

Cells were collected and fractionated as described above. After fractionation, nuclei have been lysed by sonication (20% power, 3×6s with Sonic Dismembrator Model 500, Fisher, Max Output 400W) in 0.5 ml per 4×10^7^ cells of NT-2 buffer supplemented with 1× protease inhibitors. Lysates have been cleared by centrifugation at 10,000 × g 10 min at 4°C and transferred into a clean tube, and the concentration determined by BCA assay before being adjusted to 1 mg/ml. For RNAse treatment, 1mg of nuclear lysate has been incubated at RT for 20 min with RNase A (EN0531) at 20µg/ml final and RNase T1 at 1U/µl (EN0542).

For DNaseI treatment, 1/10^th^ of DNase I buffer and 10U/ml of DNase I (invitrogen: 18047019) was added to 1 mg of extract and incubated at room temperature for 20min. Untreated or treated lysates were cleared with 30 µl of Dynabeads G per mg of extract at 4°C for 40 min and centrifuge at 10,000 × g 10 min at 4°C. For 1 mg IP, 33 µl of Dynabeads G (Invitrogen) were preincubated with 10 µg of anti-eIF4E (rabbit, MBL) or rabbit IgGs at RT for 20 min. After five washes, beads are resuspended with precleared lysate and incubated with rotation at 4°C o/n. IPs were washed one time with NT-2 buffer supplemented with 1 mg/ml Heparin and five times with 1 ml of NT-2 buffer and resuspended in 2× LB and incubated at 95°C for 10 min. Samples have been resolved by SDS–PAGE and visualized by Western Blot. Quantification of immunoprecipitation was carried out using Image Lab 6.1.0 build 7 (Bio-Rad) and plotted in PRISM 10.4.2. Differences in co-immunoprecipitation (Co-IP) signal between conditions were analyzed using two-way analysis of variance (ANOVA), with treatments (Ctrl, RNase, DNAse) and protein partner of eIF4E as independent variables. Post hoc comparisons were conducted using Dunnett’s multiple comparisons test when appropriate. Data were presented as mean ± SEM only significant differences.

### RNA immunoprecipitation

Cells were collected and fractionated as described above. After fractionation, nuclei were crosslinked with 1% PFA (Electron Microscopy) for 10 minutes at room temperature (RT). Crosslinking was quenched with 150 mM glycine (5 min, RT), and cells are centrifuged at 1000g for 5 min at 4°C and washed twice with PBS. Nuclei were resuspended and sonicated in NT2 buffer. Lysates were precleared with beads for 30 minutes at 4°C. Precleared lysates (0,5-1mg) were incubated with 5μg anti-Flag antibodies (M2, Sigma) over night at 4°C with rotation. Dynabeads G (33ul) were added per each tube. IPs were washed one time with NT-2 buffer supplemented with 1 mg/ml Heparin and additional five times with NT2 buffer with 300mM NaCl. Proteins and RNAs were eluted in Sample buffer containing 12% beta-mercaptoethanol. 15% of the elution was used for western blot and the rest was mixed with 1ml of trizol and RNAs were isolated using DirectZol kit. For RT-qPCR, RNA samples (Direct-zol RNA Miniprep Kit, Zymo Research) were reversed transcribed using Superscript Vilo reverse transcription master mix (ThermoFisher Scientific) and qPCR analyses were performed as described above.

For RNA sequencing, KAPA RNA HyperPrep Ribodepletion Kits (Roche) was used, according to the manufacturer’s instructions. Reads were trimmed and aligned as above. Differential expression analysis was carried out as above. RIP-seq data are deposited as <u>GSE345544</u>. RIP-seq targets were considered to be enriched if they had log_2_(Fold Change) > 0.58 and p_adj_ < 0.1 when compared to input. For RIP-seq, we examined three pulldowns of FLAG-eIF4E and FLAG-vector and two FLAG-NCBP2 pulldowns, compared to their respective nuclear fractionation inputs.

### Cap binding assays

Cells were fractionated as described above and nuclear fractions suspended in Cap Buffer B (50mM MOPS/KOH pH=7.4, 100mM NaCl, 50mM NaF, 2mM EDTA, 2mM EGTA, 1% NP40, 1% Na-DOC, 7mM βME, 1mM Na_3_VO_4_, and KKM protease inhibitors). 40μl m^7^GTP-Agarose beads (Creative Biomart) were washed 3 times in Cap Buffer C (50mM MOPS/KOH pH=7.4, 100mM NaCl, 50mM NaF, 0.5mM EDTA, 0.5mM EGTA, 7mM βME, 1mM Na_3_VO_4_, 0.1mM GTP, and KKM protease inhibitors). 200μg of nuclear lysate was diluted to a volume of 400μl in Cap Buffer C and the beads were added.

50μM m^7^GpppG (New England Biolabs) or equivalent volume Cap Buffer C was added, and the samples were incubated on a rotating wheel for 20 minutes at 4°C. The beads were collected via centrifugation at 500xg for 3 minutes at 4°C and washed 3x in Cap Buffer C. Cap bound proteins were eluted in 40μl 2X Laemmli loading buffer plus 7mM βME by heating at 98°C with mixing for 5 minutes. Proteins were analyzed via western blot. Quantification of blots was carried out using FIJI and plotted in PRISM using a Welch t-test.

### NEXSON Immunoprecipitation

Nuclei Extraction by sonication (NEXSON) has been firstly described in Arrigoni *et al.*^56^. Briefly, U2OS cells are resuspended by trypsinization, collected at 1200 rpm for 5 min at 4°C, and crosslinked in 1% formaldehyde (10 min, RT). Crosslinking is quenched with 125 mM glycine (5 min, RT), and cells are centrifuged at 1000g for 5 min at 4°C and washed twice with PBS. For nuclei extraction, cells are resuspended in Farnham Lab Buffer (5 mM PIPES pH 8.0, 85 mM KCl, 0.5% IGEPAL CA-630, supplemented with proteases inhibitors), sonicated with Bioruptor® Plus, 3 cycles of 15s on/30s off, 4°C, and collected by centrifugation at 1000g for 5 min at 4°C. Nuclei are washed once with PBS and resuspended in NT-2 Buffer (150mM NaCl), and sonicated (17-20%, 3×6 sec). Cells membranes and insoluble material is precipitated by centrifugation, 12000g for 10 min at 4°C. Supernatant is collected and protein concentration is measured with BCA assay. For each mg of extract, 33ul of Dynabeads G are incubated with 10µg antibody 20min at RT, washed 5times in NT-2 150 and incubated with 1mg of extract overnight at 4°C. IPs were washed one time with NT-2 buffer supplemented with 1 mg/ml Heparin and five times with 1 ml of NT-2 buffer (with 300mM NaCl). Proteins were analyzed by western blot.

### Immunofluorescence and laser-scanning confocal microscopy

U2OS cells were grown on 4 well glass slides (Millicell EZ SLIDE 4 well glass, Millipore Sigma PEZGS0416) with 5×10^5^ cells per wells for two days. After washing three times in 1× PBS (pH 7.4), cells were crosslinked in prechilled 2%PFA at 20°C for 15 min. After fixation, slides were washed three times in ice cold PBS and permeabilized in PBS + 0.2% Triton X-100 for 5 min at 20°C and blocked for 1h in Blocking solution (10% FBS and 0.1% Tween 20 in PBS). eIF4E antibody (SantaCruz A-10) was incubated overnight at 4°C in Blocking solution (1:100). After washing three times in PBS, cells were incubated with SON, SF3B1, NCBP2 or POLR2-S2P antibodies for 2 hours at room temperature. Slides were washed three times and incubated with secondary antibody (rabbit anti-RRX and mouse AF555) After washing three times in PBS, cells were mounted in antifade mounting medium with DAPI (Vector Laboratories, H2000).

Analysis was carried out using a laser-scanning confocal microscope (Leica TCS SP8 DLS, LightSheet), 63× oil objective and numerical aperture of 1. Channels were detected separately, with no crosstalk observed. Confocal micrographs represent single sections through the plane of the cell. Images were obtained from LAS-X software (Leica Microsystems) and displayed using Adobe Photoshop CS6 (Adobe).

For quantification of co-localization images were opened with Imaris and the background was determined using control images of cells labeled with rabbit and mouse IgGs, followed by incubation with secondary antibodies (anti-mouse RRX and anti-rabbit AF488). The threshold was set to 50 for both colors. For quantification, multi-cell images were cropped to obtain at least 20 individual nuclei per condition. A DAPI mask was applied to specifically quantify the signal within the nuclear region of the cells. A colocalization channel was generated, and statistics were extracted from it. Mander’s coefficient was used to assess the degree of co-localization between the two signals.

The percent co-localization was based on the ROI volume colocalized.

### Data analysis and visualization

Analysis and visualization of the differential gene expression and alternate splicing data was done in R version 4.4.2 using the *tidyverse* suite of packages for data manipulation, *biomaRt* and *BSgenome* to retrieve genomic data, *pheatmap* for making heatmaps, *eulerr* for making Euler diagrams, and *ggplot2* for all other plots. The web-based version of metascape was used (https://metascape.org/gp/index.html#/main/step1) for gene ontology analysis employing default settings and accessed in June-July 2025 for the overexpression datasets and June-August 2026 for the knockdown and RIP-seq datasets^85^. Analysis and visualization of qPCR was performed in PRISM 10.4.2.

## Supporting information

Apendix 1

## Acknowledgements

We thank Jason Tien and Xiaopei Zhang for technical assistance. We are grateful to the genomics and confocal microscopy platforms at University of Montreal and Northwestern University for their technical expertise. We are grateful to Jack Keene for 3A2 anti-HuR antibody. This work was supported by the NIH, CME was supported by NIH/NCI training grant T32 CA070085, and KLBB by NIH/NCI grants RO1 CA98571 and RO1 CA80728, the Canadian Institute of Health Research (PJT 159785), and the Canada Research Chair programme.

## Disclosing interest statements

JCM, CME, BCK, GV, NS, PG, AWBC and KLBB have no disclosure or competing interests.

---

**Figure S1:**
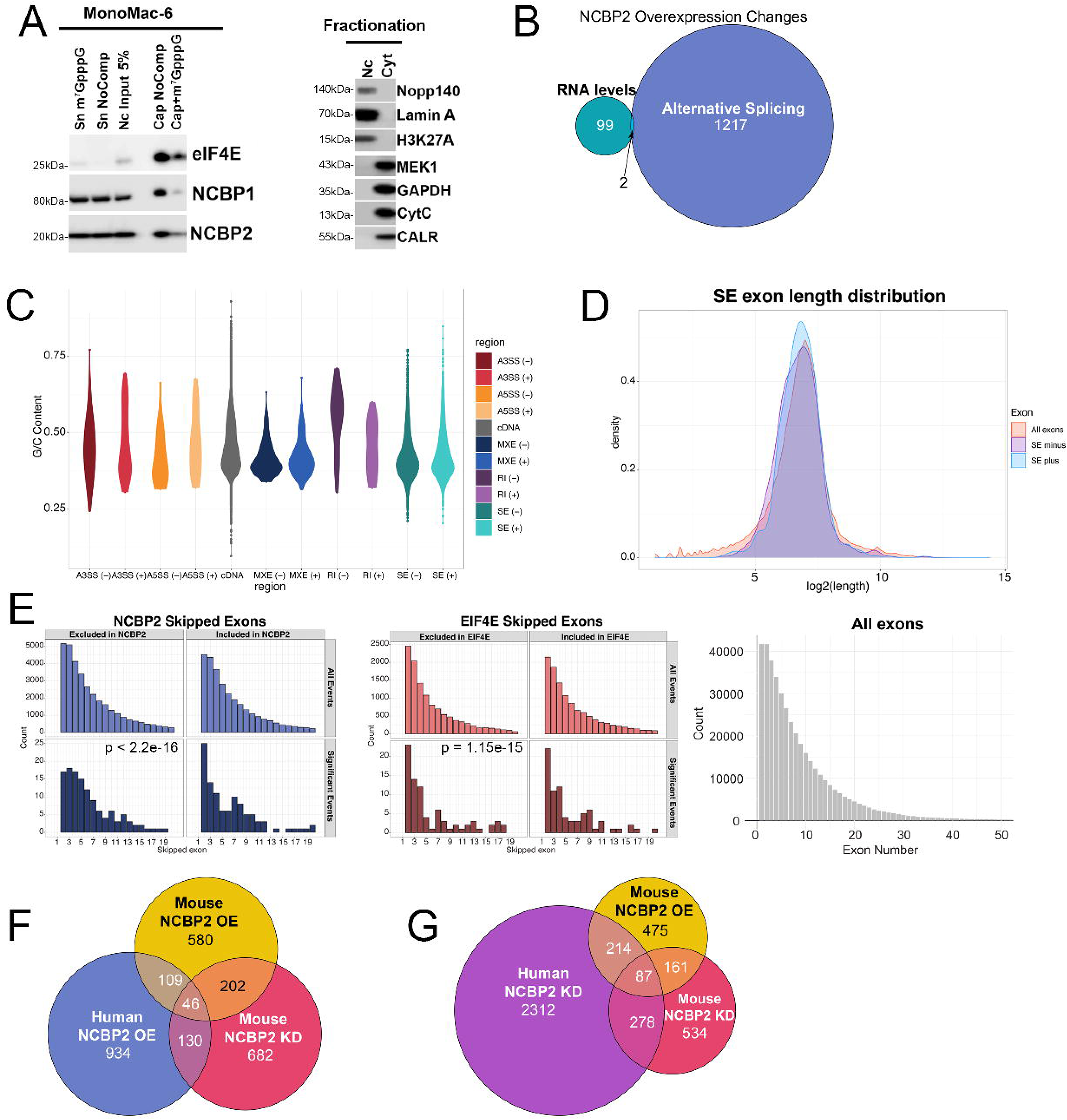
eIF4E and NCBP2 splicing targets have shared characteristics. **A)** *Left* Western blot showing cap chromatography of nuclear lysate in MonoMac-6 cells. *Right* Fractionation controls for the MonoMac-6 cap chromatography. **B)** Euler plot showing the overlap between transcripts undergoing differential expression (green) and alternative splicing (blue) following NCBP2 overexpression. **C)** Violin plot showing the GC content of transcripts with A3SS (red), A5SS (orange), MXE (blue), RI (purple), SE (teal), or all transcripts (grey) in NCBP2 overexpression. Lighter shades indicate events with more inclusion in NCBP2 (+). **D)** Density plot showing the length of all exons (red) or skipped exons that are more (blue) or less (purple) included in NCBP2 overexpression. **E)** Position of skipped exons in following overexpression of NCBP2 (blue, left) or eIF4E (red, middle). All detected skipped exons are shown on the top of each plot and only significant events are shown on the bottom. The similarity of the distribution of significant skipped exons were compared to the distribution of all exons w transcript support level 1 transcripts (right) with a χ^2^ goodness of fit test with the p-value indicated. **F)** Euler plot showing the overlap between transcripts undergoing alternative splicing as a result of NCBP2 overexpression in U2OS cells (blue), NCBP2 overexpression in mouse cardiomyocyte cells (yellow), or NCBP2 knockdown in mouse cardiomyocyte cells (pink). **F)** Euler plot showing the overlap between transcripts undergoing alternative splicing as a result of NCBP2 knockdown in U2OS cells (purple), NCBP2 overexpression in mouse cardiomyocyte cells (yellow), or NCBP2 knockdown in mouse cardiomyocyte cells (pink).

**Figure S2:**
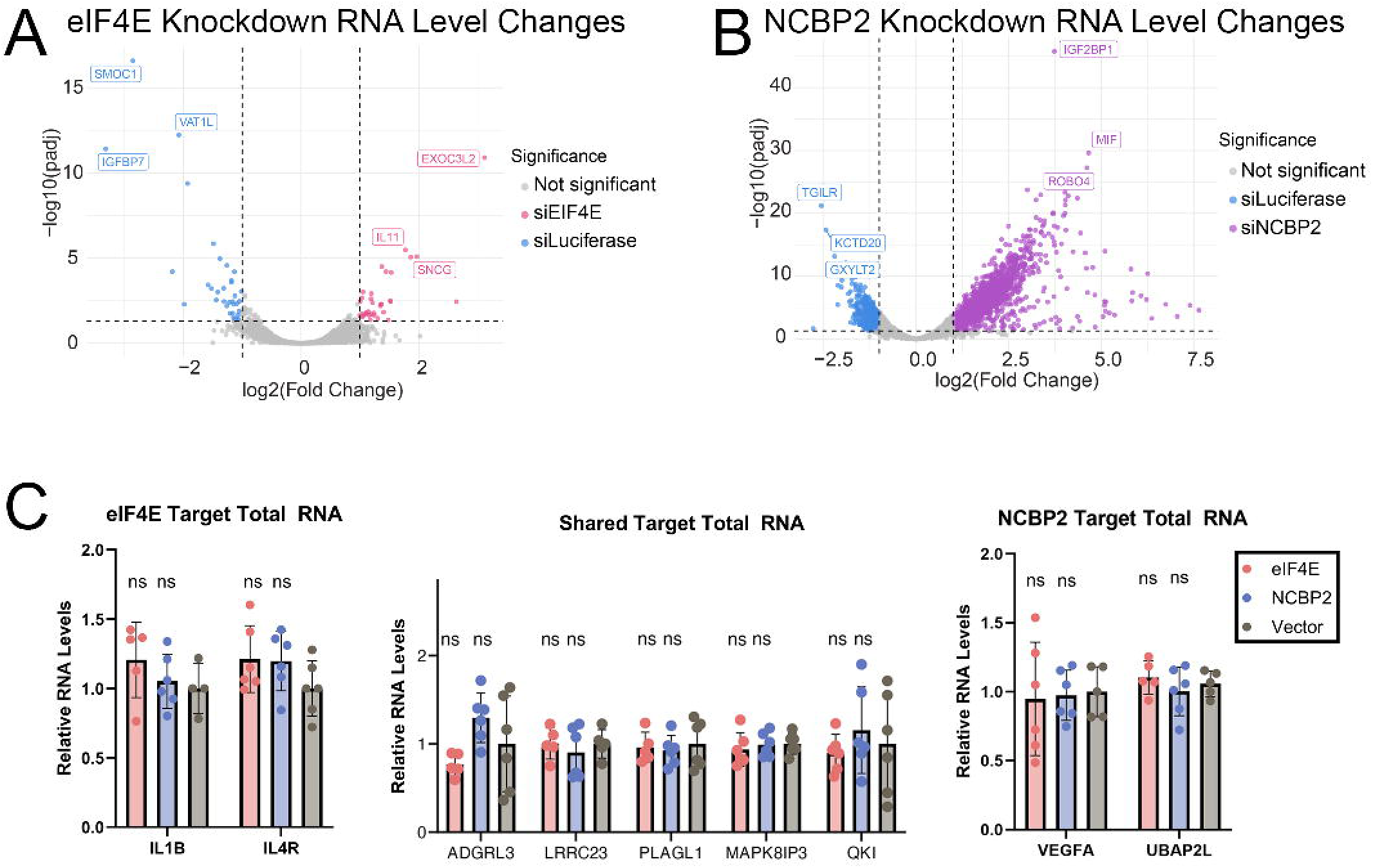
eIF4E and NCBP2 knockdown cause different transcriptomic changes. **A)** Volcano plot showing the upregulated (pink) and downregulated (blue) transcripts in eIF4E knockdown. Vertical dashed lines indicate 2-fold change and horizontal black line indicates padj = 0.05. **B)** Volcano plot showing the upregulated (purple) and downregulated (blue) transcripts in NCBP2 knockdown. Vertical dashed lines indicate 2-fold change and horizontal black line indicates padj = 0.05. **C)** Quantitative PCR showing the total RNA levels following NCBP2 (blue) or eIF4E (red) overexpression. * = p < 0.05, ns = non-significant.

**Figure S3:**
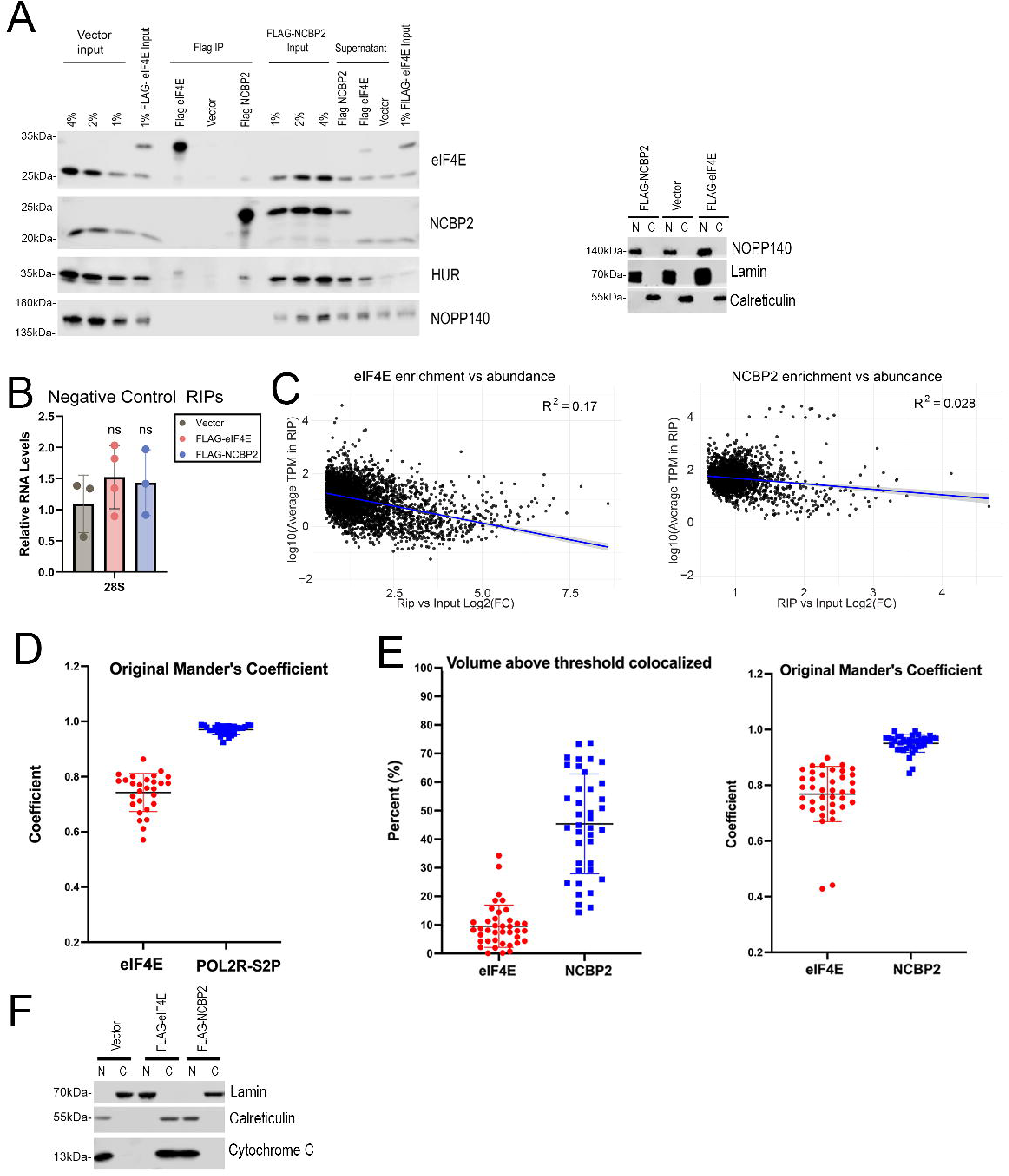
eIF4E and NCBP2 have substantial segregation in the nucleus. **A)** *Left* Western blot showing the quality of the FLAG-IP used for FLAG-eIF4E and FLAG-NCBP2 RTqPCR and RIP-seq. *Right* Fractionation control for RIP nuclear fractionation. **B)** Negative controls showing 28S rRNA is not pulled down in RIPs from nuclear lysates: FLAG-vector (grey), FLAG-eIF4E (red), or FLAG-NCBP2 (blue). ns = non-significant **C)** Scatter plots showing no correlation between enrichment and average transcript abundance in FLAG-eIF4E (left) and FLAG-NCBP2 (right) RIP-seq. **D)** The Manders coefficient for Figure 5E. Each point represents a single nuclei. Data were collected as Z-stacks for >25 cells and processed in PRISM. **E)** Quantitation of Figure 6A colocalization using Imaris. Percentage colocalization over the cell volume (left) and the Manders coefficient (right) are shown. Each point represents a single nuclei. Data were collected as Z-stacks for >25 cells and processed in PRISM. **F)** Fractionation control for FLAG-eIF4E and FLAG-NCBP2 immunoprecipitations shown in Figure 6E

**Figure S4:**
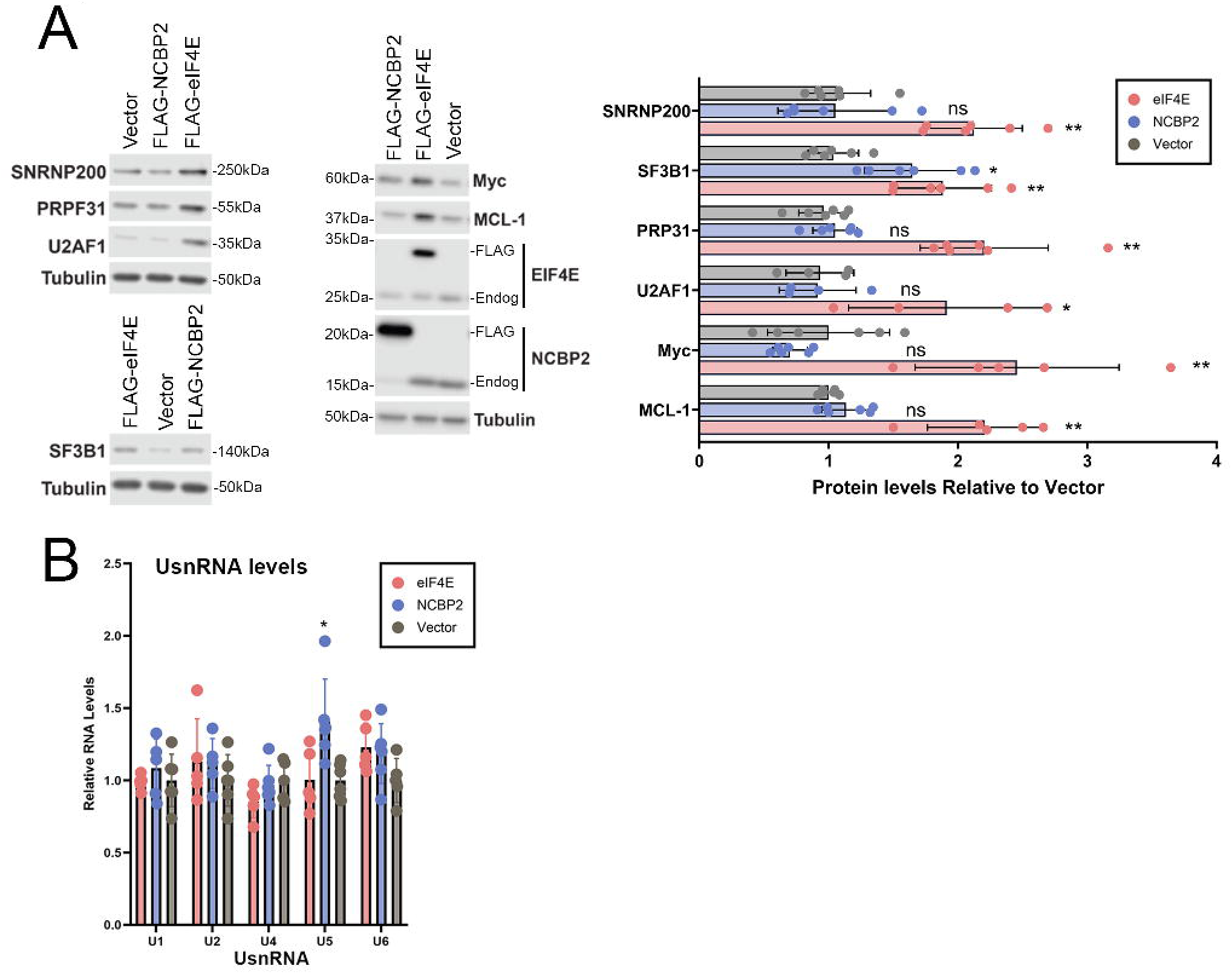
eIF4E and NCBP2 overexpression have distinct effects on splice factor production. **A)** *Left* Western blot showing the impact of eIF4E or NCBP2 overexpression on production of each other and selected splice factors. Representative blots probed as indicated. Endog refers to endogenous NCBP2 or eIF4E and FLAG, the overexpressed form. *Right* Quantitation of 3-6 biological replicates for each condition are shown. **B)** RT-qPCR analysis of UsnRNA levels in U2OS cells overexpressing eIF4E or NCBP2. For all panels, means, standard deviations and p-values relative to vector (Welch t-test) were calculated in PRISM. * = p-value < 0.05, ** = p-value < 0.01, n.s. = non-significant.

## References

1. Borden, K., Culjkovic-Kraljacic, B., and Cowling, V.H. (2021). To cap it all off, again: dynamic capping and recapping of coding and non-coding RNAs to control transcript fate and biological activity. Cell Cycle 20, 1347–1360. 10.1080/15384101.2021.1930929.

2. Cowling, V.H. (2009). Regulation of mRNA cap methylation. The Biochemical journal 425, 295–302. 10.1042/BJ20091352.

3. Schoenberg, D.R., and Maquat, L.E. (2009). Re-capping the message. Trends in biochemical sciences 34, 435–442. 10.1016/j.tibs.2009.05.003.

4. Rasmussen, E.B., and Lis, J.T. (1993). In vivo transcriptional pausing and cap formation on three Drosophila heat shock genes. Proc Natl Acad Sci U S A 90, 7923–7927. 10.1073/pnas.90.17.7923.

5. Rambout, X., and Maquat, L.E. (2020). The nuclear cap-binding complex as choreographer of gene transcription and pre-mRNA processing. Genes Dev 34, 1113–1127. 10.1101/gad.339986.120.

6. Lykke-Andersen, S., Rouviere, J.O., and Jensen, T.H. (2021). ARS2/SRRT: at the nexus of RNA polymerase II transcription, transcript maturation and quality control. Biochem Soc Trans 49, 1325–1336. 10.1042/BST20201008.

7. Neugebauer, K.M. (2002). On the importance of being co-transcriptional. J Cell Sci 115, 3865–3871. 10.1242/jcs.00073.

8. Kentsis, A., and Borden, K.L. (2000). Construction of macromolecular assemblages in eukaryotic processes and their role in human disease: linking RINGs together. Curr Protein Pept Sci 1, 49–73. 10.2174/1389203003381478.

9. Giacometti, S., Benbahouche, N.E.H., Domanski, M., Robert, M.C., Meola, N., Lubas, M., Bukenborg, J., Andersen, J.S., Schulze, W.M., Verheggen, C., et al. (2017). Mutually Exclusive CBC-Containing Complexes Contribute to RNA Fate. Cell Rep 18, 2635–2650. 10.1016/j.celrep.2017.02.046.

10. Bush, M.S., Hutchins, A.P., Jones, A.M., Naldrett, M.J., Jarmolowski, A., Lloyd, C.W., and Doonan, J.H. (2009). Selective recruitment of proteins to 5’ cap complexes during the growth cycle in Arabidopsis. Plant J 59, 400–412. 10.1111/j.1365-313X.2009.03882.x.

11. Ishigaki, Y., Li, X., Serin, G., and Maquat, L.E. (2001). Evidence for a pioneer round of mRNA translation: mRNAs subject to nonsense-mediated decay in mammalian cells are bound by CBP80 and CBP20. Cell 106, 607–617. 10.1016/s0092-8674(01)00475-5.

12. Lejeune, F., Ishigaki, Y., Li, X., and Maquat, L.E. (2002). The exon junction complex is detected on CBP80-bound but not eIF4E-bound mRNA in mammalian cells: dynamics of mRNP remodeling. EMBO J 21, 3536–3545. 10.1093/emboj/cdf345.

13. Lejbkowicz, F., Goyer, C., Darveau, A., Neron, S., Lemieux, R., and Sonenberg, N. (1992). A fraction of the mRNA 5’ cap-binding protein, eukaryotic initiation factor 4E, localizes to the nucleus. Proc Natl Acad Sci U S A 89, 9612–9616. 10.1073/pnas.89.20.9612.

14. Lang, V., Zanchin, N.I., Lunsdorf, H., Tuite, M., and McCarthy, J.E. (1994). Initiation factor eIF-4E of Saccharomyces cerevisiae. Distribution within the cell, binding to mRNA, and consequences of its overproduction. J Biol Chem 269, 6117–6123.

15. Cohen, N., Sharma, M., Kentsis, A., Perez, J.M., Strudwick, S., and Borden, K.L. (2001). PML RING suppresses oncogenic transformation by reducing the affinity of eIF4E for mRNA. EMBO J 20, 4547–4559. 10.1093/emboj/20.16.4547.

16. Culjkovic-Kraljacic, B., Fernando, T.M., Marullo, R., Calvo-Vidal, N., Verma, A., Yang, S., Tabbo, F., Gaudiano, M., Zahreddine, H., Goldstein, R.L., et al. (2016). Combinatorial targeting of nuclear export and translation of RNA inhibits aggressive B-cell lymphomas. Blood 127, 858–868. 10.1182/blood-2015-05-645069.

17. Culjkovic, B., Topisirovic, I., Skrabanek, L., Ruiz-Gutierrez, M., and Borden, K.L. (2005). eIF4E promotes nuclear export of cyclin D1 mRNAs via an element in the 3’UTR. J Cell Biol 169, 245–256. 10.1083/jcb.200501019.

18. Dostie, J., Lejbkowicz, F., and Sonenberg, N. (2000). Nuclear eukaryotic initiation factor 4E (eIF4E) colocalizes with splicing factors in speckles. J Cell Biol 148, 239–247. 10.1083/jcb.148.2.239.

19. Gebhardt, A., Habjan, M., Benda, C., Meiler, A., Haas, D.A., Hein, M.Y., Mann, A., Mann, M., Habermann, B., and Pichlmair, A. (2015). mRNA export through an additional cap-binding complex consisting of NCBP1 and NCBP3. Nat Commun 6, 8192. 10.1038/ncomms9192.

20. Ohno, M., Kataoka, N., and Shimura, Y. (1990). A nuclear cap binding protein from HeLa cells. Nucleic Acids Res 18, 6989–6995. 10.1093/nar/18.23.6989.

21. Embree, C.M., and Borden, K.L.B. (2026). Escorting mRNAs - the multifunctional and divergent roles of cap-chaperones in post-transcriptional gene expression. J Cell Sci 139. 10.1242/jcs.264780.

22. Schulze, W.M., Stein, F., Rettel, M., Nanao, M., and Cusack, S. (2018). Structural analysis of human ARS2 as a platform for co-transcriptional RNA sorting. Nat Commun 9, 1701. 10.1038/s41467-018-04142-7.

23. Kentsis, A., Topisirovic, I., Culjkovic, B., Shao, L., and Borden, K.L. (2004). Ribavirin suppresses eIF4E-mediated oncogenic transformation by physical mimicry of the 7-methyl guanosine mRNA cap. Proc Natl Acad Sci U S A 101, 18105–18110. 10.1073/pnas.0406927102.

24. Dubiez, E., Pellegrini, E., Finderup Brask, M., Garland, W., Foucher, A.E., Huard, K., Heick Jensen, T., Cusack, S., and Kadlec, J. (2024). Structural basis for competitive binding of productive and degradative co-transcriptional effectors to the nuclear cap-binding complex. Cell Rep 43, 113639. 10.1016/j.celrep.2023.113639.

25. Mars, J.C., Culjkovic-Kraljacic, B., and Borden, K.L.B. (2024). eIF4E orchestrates mRNA processing, RNA export and translation to modify specific protein production. Nucleus 15, 2360196. 10.1080/19491034.2024.2360196.

26. Davis, M.R., Delaleau, M., and Borden, K.L.B. (2019). Nuclear eIF4E Stimulates 3’-End Cleavage of Target RNAs. Cell Rep 27, 1397–1408 e1394. 10.1016/j.celrep.2019.04.008.

27. Culjkovic-Kraljacic, B., Skrabanek, L., Revuelta, M.V., Gasiorek, J., Cowling, V.H., Cerchietti, L., and Borden, K.L.B. (2020). The eukaryotic translation initiation factor eIF4E elevates steady-state m(7)G capping of coding and noncoding transcripts. Proc Natl Acad Sci U S A 117, 26773–26783. 10.1073/pnas.2002360117.

28. Culjkovic, B., Tan, K., Orolicki, S., Amri, A., Meloche, S., and Borden, K.L. (2008). The eIF4E RNA regulon promotes the Akt signaling pathway. J Cell Biol 181, 51–63. 10.1083/jcb.200707018.

29. Culjkovic, B., Topisirovic, I., Skrabanek, L., Ruiz-Gutierrez, M., and Borden, K.L. (2006). eIF4E is a central node of an RNA regulon that governs cellular proliferation. J Cell Biol 175, 415–426. 10.1083/jcb.200607020.

30. Volpon, L., Culjkovic-Kraljacic, B., Sohn, H.S., Blanchet-Cohen, A., Osborne, M.J., and Borden, K.L.B. (2017). A biochemical framework for eIF4E-dependent mRNA export and nuclear recycling of the export machinery. RNA 23, 927–937. 10.1261/rna.060137.116.

31. Topisirovic, I., Siddiqui, N., Lapointe, V.L., Trost, M., Thibault, P., Bangeranye, C., Pinol-Roma, S., and Borden, K.L. (2009). Molecular dissection of the eukaryotic initiation factor 4E (eIF4E) export-competent RNP. EMBO J 28, 1087–1098. 10.1038/emboj.2009.53.

32. Osborne, M.J., Volpon, L., Memarpoor-Yazdi, M., Pillay, S., Thambipillai, A., Czarnota, S., Culjkovic-Kraljacic, B., Trahan, C., Oeffinger, M., Cowling, V.H., and Borden, K.L.B. (2022). Identification and Characterization of the Interaction Between the Methyl-7-Guanosine Cap Maturation Enzyme RNMT and the Cap-Binding Protein eIF4E. J Mol Biol 434, 167451. 10.1016/j.jmb.2022.167451.

33. Ghram, M., Morris, G., Culjkovic-Kraljacic, B., Mars, J.C., Gendron, P., Skrabanek, L., Revuelta, M.V., Cerchietti, L., Guzman, M.L., and Borden, K.L.B. (2023). The eukaryotic translation initiation factor eIF4E reprograms alternative splicing. EMBO J 42, e110496. 10.15252/embj.2021110496.

34. Lenasi, T., Peterlin, B.M., and Barboric, M. (2011). Cap-binding protein complex links pre-mRNA capping to transcription elongation and alternative splicing through positive transcription elongation factor b (P-TEFb). J Biol Chem 286, 22758–22768. 10.1074/jbc.M111.235077.

35. Izaurralde, E., Lewis, J., McGuigan, C., Jankowska, M., Darzynkiewicz, E., and Mattaj, I.W. (1994). A nuclear cap binding protein complex involved in pre-mRNA splicing. Cell 78, 657–668. 10.1016/0092-8674(94)90530-4.

36. Colot, H.V., Stutz, F., and Rosbash, M. (1996). The yeast splicing factor Mud13p is a commitment complex component and corresponds to CBP20, the small subunit of the nuclear cap-binding complex. Genes Dev 10, 1699–1708. 10.1101/gad.10.13.1699.

37. Li, Z., Cao, C., Zhao, Q., Li, D., Han, Y., Zhang, M., Mao, L., Zhou, B., and Wang, L. (2025). RNA splicing controls organ-wide maturation of postnatal heart in mice. Dev Cell 60, 236–252 e238. 10.1016/j.devcel.2024.09.018.

38. Pabis, M., Neufeld, N., Steiner, M.C., Bojic, T., Shav-Tal, Y., and Neugebauer, K.M. (2013). The nuclear cap-binding complex interacts with the U4/U6.U5 tri-snRNP and promotes spliceosome assembly in mammalian cells. RNA 19, 1054–1063. 10.1261/rna.037069.112.

39. Fortes, P., Inada, T., Preiss, T., Hentze, M.W., Mattaj, I.W., and Sachs, A.B. (2000). The yeast nuclear cap binding complex can interact with translation factor eIF4G and mediate translation initiation. Molecular cell 6, 191–196.

40. Assouline, S., Culjkovic, B., Cocolakis, E., Rousseau, C., Beslu, N., Amri, A., Caplan, S., Leber, B., Roy, D.C., Miller, W.H., Jr., and Borden, K.L. (2009). Molecular targeting of the oncogene eIF4E in acute myeloid leukemia (AML): a proof-of-principle clinical trial with ribavirin. Blood 114, 257–260. 10.1182/blood-2009-02-205153.

41. Assouline, S., Culjkovic-Kraljacic, B., Bergeron, J., Caplan, S., Cocolakis, E., Lambert, C., Lau, C.J., Zahreddine, H.A., Miller, W.H., Jr., and Borden, K.L. (2015). A phase I trial of ribavirin and low-dose cytarabine for the treatment of relapsed and refractory acute myeloid leukemia with elevated eIF4E. Haematologica 100, e7–9. 10.3324/haematol.2014.111245.

42. Assouline, S., Gasiorek, J., Bergeron, J., Lambert, C., Culjkovic-Kraljacic, B., Cocolakis, E., Zakaria, C., Szlachtycz, D., Yee, K., and Borden, K.L.B. (2023). Molecular targeting of the UDP-glucuronosyltransferase enzymes in high-eukaryotic translation initiation factor 4E refractory/relapsed acute myeloid leukemia patients: a randomized phase II trial of vismodegib, ribavirin with or without decitabine. Haematologica. 10.3324/haematol.2023.282791.

43. Li, S., Wang, Y., Yang, X., Li, M., Li, G., Song, Q., and Liu, J. (2025). NCBP2 predicts the prognosis and the immunotherapy response of cancers: a pan-cancer analysis. PeerJ 13, e19050. 10.7717/peerj.19050.

44. Wang, J., Guo, T., Zhang, X., Guo, J., Meng, X., Yan, S., Wang, Y., Xiao, Y., Xu, W., Wei, X., et al. (2024). Comprehensive investigation in oncogenic functions and immunological roles of NCBP2 and its validation in prostate cancer. Transl Oncol 47, 102049. 10.1016/j.tranon.2024.102049.

45. Liu, L., Lu, W., Miao, S., Yu, Y., Zhang, X.B., Ye, S., Wang, X., and Liu, L. (2026). NCBP2 drives colorectal cancer growth and metastasis through LIPG-mediated lipid droplet accumulation. Commun Biol 9. 10.1038/s42003-026-09903-5.

46. Culjkovic-Kraljacic B, S.L., Revuelta M, Gasiorek J, Cowling VH, Cerchietti L, Borden KL. (2020). Gene Expression Omnibus GSE 158728 (https://www.ncbi.nlm.nih.gov/geo/query/acc.cgi?acc=GSE158728<u>).[DATASET</u>].

47. Hariri, F., Arguello, M., Volpon, L., Culjkovic-Kraljacic, B., Nielsen, T.H., Hiscott, J., Mann, K.K., and Borden, K.L. (2013). The eukaryotic translation initiation factor eIF4E is a direct transcriptional target of NF-kappaB and is aberrantly regulated in acute myeloid leukemia. Leukemia 27, 2047–2055. 10.1038/leu.2013.73.

48. Shen, S., Park, J.W., Huang, J., Dittmar, K.A., Lu, Z.X., Zhou, Q., Carstens, R.P., and Xing, Y. (2012). MATS: a Bayesian framework for flexible detection of differential alternative splicing from RNA-Seq data. Nucleic Acids Res 40, e61. 10.1093/nar/gkr1291.

49. Cordiner, R.A., Dou, Y., Thomsen, R., Bugai, A., Granneman, S., and Heick Jensen, T. (2023). Temporal-iCLIP captures co-transcriptional RNA-protein interactions. Nat Commun 14, 696. 10.1038/s41467-023-36345-y.

50. Dong, K., Wang, R., Wang, X., Lin, F., Shen, J.J., Gao, P., and Zhang, H.Z. (2009). Tumor-specific RNAi targeting eIF4E suppresses tumor growth, induces apoptosis and enhances cisplatin cytotoxicity in human breast carcinoma cells. Breast Cancer Res Treat 113, 443–456. 10.1007/s10549-008-9956-x.

51. Cho, Y., Lee, S.E., Oh, J., Jang, D., Shin, S., Lee, S.J., Kim, J., Yang, Y., Kim, D., Jung, H.R., et al. (2025). Comprehensive profiling of RNA modification-related genes identifies RNA m(7)G binding protein CBP20 as a therapeutic target for tumor growth inhibition. Exp Mol Med 57, 1978–1995. 10.1038/s12276-025-01531-z.

52. Cherla, R.P., Lee, S.Y., Mees, P.L., and Tesh, V.L. (2006). Shiga toxin 1-induced cytokine production is mediated by MAP kinase pathways and translation initiation factor eIF4E in the macrophage-like THP-1 cell line. J Leukoc Biol 79, 397–407. 10.1189/jlb.0605313.

53. Medford, A.R., Douglas, S.K., Godinho, S.I., Uppington, K.M., Armstrong, L., Gillespie, K.M., van Zyl, B., Tetley, T.D., Ibrahim, N.B., and Millar, A.B. (2009). Vascular Endothelial Growth Factor (VEGF) isoform expression and activity in human and murine lung injury. Respir Res 10, 27. 10.1186/1465-9921-10-27.

54. Phatnani, H.P., and Greenleaf, A.L. (2006). Phosphorylation and functions of the RNA polymerase II CTD. Genes Dev 20, 2922–2936. 10.1101/gad.1477006.

55. Komarnitsky, P., Cho, E.J., and Buratowski, S. (2000). Different phosphorylated forms of RNA polymerase II and associated mRNA processing factors during transcription. Genes Dev 14, 2452–2460. 10.1101/gad.824700.

56. Arrigoni, L., Richter, A.S., Betancourt, E., Bruder, K., Diehl, S., Manke, T., and Bonisch, U. (2016). Standardizing chromatin research: a simple and universal method for ChIP-seq. Nucleic Acids Res 44, e67. 10.1093/nar/gkv1495.

57. Beacon, T.H., Delcuve, G.P., Lopez, C., Nardocci, G., Kovalchuk, I., van Wijnen, A.J., and Davie, J.R. (2021). The dynamic broad epigenetic (H3K4me3, H3K27ac) domain as a mark of essential genes. Clin Epigenetics 13, 138. 10.1186/s13148-021-01126-1.

58. Palangat, M., Renner, D.B., Price, D.H., and Landick, R. (2005). A negative elongation factor for human RNA polymerase II inhibits the anti-arrest transcript-cleavage factor TFIIS. Proc Natl Acad Sci U S A 102, 15036–15041. 10.1073/pnas.0409405102.

59. Aoi, Y., Smith, E.R., Shah, A.P., Rendleman, E.J., Marshall, S.A., Woodfin, A.R., Chen, F.X., Shiekhattar, R., and Shilatifard, A. (2020). NELF Regulates a Promoter-Proximal Step Distinct from RNA Pol II Pause-Release. Molecular cell 78, 261–274 e265. 10.1016/j.molcel.2020.02.014.

60. Fong, N., Ohman, M., and Bentley, D.L. (2009). Fast ribozyme cleavage releases transcripts from RNA polymerase II and aborts co-transcriptional pre-mRNA processing. Nat Struct Mol Biol 16, 916–922. 10.1038/nsmb.1652.

61. Topisirovic, I., Capili, A.D., and Borden, K.L. (2002). Gamma interferon and cadmium treatments modulate eukaryotic initiation factor 4E-dependent mRNA transport of cyclin D1 in a PML-dependent manner. Mol Cell Biol 22, 6183–6198. 10.1128/MCB.22.17.6183-6198.2002.

62. Xie, J., Mo, T., Li, R., Zhang, H., Liang, G., Ma, T., Chen, J., Xie, H., Wen, X., Hu, T., et al. (2023). The m(7)G Reader NCBP2 Promotes Pancreatic Cancer Progression by Upregulating MAPK/ERK Signaling. Cancers (Basel) 15. 10.3390/cancers15225454.

63. Zhu, W., Yang, W., Sun, G., and Huang, J. (2025). RNA-binding protein quaking: a multifunctional regulator in tumour progression. Ann Med 57, 2443046. 10.1080/07853890.2024.2443046.

64. Teplova, M., Hafner, M., Teplov, D., Essig, K., Tuschl, T., and Patel, D.J. (2013). Structure-function studies of STAR family Quaking proteins bound to their in vivo RNA target sites. Genes Dev 27, 928–940. 10.1101/gad.216531.113.

65. Chen, T., and Richard, S. (1998). Structure-function analysis of Qk1: a lethal point mutation in mouse quaking prevents homodimerization. Mol Cell Biol 18, 4863–4871. 10.1128/MCB.18.8.4863.

66. Justice, M.J., and Bode, V.C. (1988). Three ENU-induced alleles of the murine quaking locus are recessive embryonic lethal mutations. Genet Res 51, 95–102. 10.1017/s0016672300024101.

67. Hu, M., Yuan, X., Liu, Y., Tang, S., Miao, J., Zhou, Q., and Chen, S. (2017). IL-1beta-induced NF-kappaB activation down-regulates miR-506 expression to promotes osteosarcoma cell growth through JAG1. Biomed Pharmacother 95, 1147–1155. 10.1016/j.biopha.2017.08.120.

68. Liu, B., Zhou, Y., Chen, X., and Peng, D. (2017). IL-1beta-mediated NF-kappaB signaling augments the osteosarcoma cell growth through modulating miR-376c/TGFA axis. Pharmazie 72, 419–424. 10.1691/ph.2017.6888.

69. Peach, C.J., Mignone, V.W., Arruda, M.A., Alcobia, D.C., Hill, S.J., Kilpatrick, L.E., and Woolard, J. (2018). Molecular Pharmacology of VEGF-A Isoforms: Binding and Signalling at VEGFR2. Int J Mol Sci 19. 10.3390/ijms19041264.

70. Sato, T., Ishikawa, M., Mochizuki, M., Ohta, M., Ohkura, M., Nakai, J., Takamatsu, N., and Yoshioka, K. (2015). JSAP1/JIP3 and JLP regulate kinesin-1-dependent axonal transport to prevent neuronal degeneration. Cell Death Differ 22, 1260–1274. 10.1038/cdd.2014.207.

71. Qiu, J., Cheng, F., and Pintel, D. (2007). Distance-dependent processing of adeno-associated virus type 5 RNA is controlled by 5’ exon definition. J Virol 81, 7974–7984. 10.1128/JVI.00714-07.

72. Narita, T., Yung, T.M., Yamamoto, J., Tsuboi, Y., Tanabe, H., Tanaka, K., Yamaguchi, Y., and Handa, H. (2007). NELF interacts with CBC and participates in 3’ end processing of replication-dependent histone mRNAs. Molecular cell 26, 349–365. 10.1016/j.molcel.2007.04.011.

73. Lewis, J.D., Izaurralde, E., Jarmolowski, A., McGuigan, C., and Mattaj, I.W. (1996). A nuclear cap-binding complex facilitates association of U1 snRNP with the cap-proximal 5’ splice site. Genes Dev 10, 1683–1698. 10.1101/gad.10.13.1683.

74. Gornemann, J., Kotovic, K.M., Hujer, K., and Neugebauer, K.M. (2005). Cotranscriptional spliceosome assembly occurs in a stepwise fashion and requires the cap binding complex. Molecular cell 19, 53–63. 10.1016/j.molcel.2005.05.007.

75. Lazaris-Karatzas, A., Montine, K.S., and Sonenberg, N. (1990). Malignant transformation by a eukaryotic initiation factor subunit that binds to mRNA 5’ cap. Nature 345, 544–547.

76. De Benedetti, A., and Rhoads, R.E. (1990). Overexpression of eukaryotic protein synthesis initiation factor 4E in HeLa cells results in aberrant growth and morphology. Proc Natl Acad Sci U S A 87, 8212–8216. 10.1073/pnas.87.21.8212.

77. Arora, R., Haynes, L., Kumar, M., McNeil, R., Ashkani, J., Nakoneshny, S.C., Matthews, T.W., Chandarana, S., Hart, R.D., Jones, S.J.M., et al. (2023). NCBP2 and TFRC are novel prognostic biomarkers in oral squamous cell carcinoma. Cancer Gene Ther 30, 752–765. 10.1038/s41417-022-00578-8.

78. Bu, H., Cao, T., Li, X., Guo, Y., Guo, J., Wang, Y., Sun, Y., and Wang, D. (2022). Diagnostic and prognostic potential of the novel biomarker nuclear cap binding protein subunit 2 (NCBP2) in colon adenocarcinoma. J Gastrointest Oncol 13, 1782–1792. 10.21037/jgo-22-665.

79. Sung, H., Kang, S.H., Bae, Y.J., Hong, J.T., Chung, Y.B., Lee, C.K., and Song, S. (2006). PCR-based detection of Mycoplasma species. J Microbiol 44, 42–49.

80. Zahreddine, H.A., Culjkovic-Kraljacic, B., Emond, A., Pettersson, F., Midura, R., Lauer, M., Del Rincon, S., Cali, V., Assouline, S., Miller, W.H., et al. (2017). The eukaryotic translation initiation factor eIF4E harnesses hyaluronan production to drive its malignant activity. Elife 6. 10.7554/eLife.29830.

81. Bolger, T.A., Folkmann, A.W., Tran, E.J., and Wente, S.R. (2008). The mRNA export factor Gle1 and inositol hexakisphosphate regulate distinct stages of translation. Cell 134, 624–633. 10.1016/j.cell.2008.06.027.

82. Dobin, A., Davis, C.A., Schlesinger, F., Drenkow, J., Zaleski, C., Jha, S., Batut, P., Chaisson, M., and Gingeras, T.R. (2013). STAR: ultrafast universal RNA-seq aligner. Bioinformatics 29, 15–21. 10.1093/bioinformatics/bts635.

83. Love, M.I., Huber, W., and Anders, S. (2014). Moderated estimation of fold change and dispersion for RNA-seq data with DESeq2. Genome Biol 15, 550. 10.1186/s13059-014-0550-8.

84. Wang, Y., Xie, Z., Kutschera, E., Adams, J.I., Kadash-Edmondson, K.E., and Xing, Y. (2024). rMATS-turbo: an efficient and flexible computational tool for alternative splicing analysis of large-scale RNA-seq data. Nat Protoc 19, 1083–1104. 10.1038/s41596-023-00944-2.

85. Zhou, Y., Zhou, B., Pache, L., Chang, M., Khodabakhshi, A.H., Tanaseichuk, O., Benner, C., and Chanda, S.K. (2019). Metascape provides a biologist-oriented resource for the analysis of systems-level datasets. Nat Commun 10, 1523. 10.1038/s41467-019-09234-6.

