## Supplementary material for "Nuclear RNA cap-chaperones eIF4E and NCBP2 influence distinct mRNA fates": Table 1

**Table 1. Biological Consequences of Selected Cap-Chaperone Regulated Splicing Events**

| **Target mRNA** | **Splicing Event (upon overexpression)** | **RIP Fold Enrichment — Splice-Event-Specific Primers** | | **RIP Fold Enrichment — Universal Primers (all isoforms)** | | **Biological Consequence** |
| --- | --- | --- | --- | --- | --- | --- |
|  |  | **eIF4E** | **NCBP2** | **eIF4E** | **NCBP2** |  |
| **eIF4E-sensitive targets** | | | | | | |
| **IL1B** | **1st intron retention (IR)**  *Reduced intron retention upon eIF4E overexpression* | **3.9 ± 0.86**  **p < 0.005** | 2.11 ± 0.86  P = 0.13 | **19.19 ± 5.05**  **P < 0.05** | **11.79 ± 1.05**  **P < 0.005** | Reduced intron retention will increase IL1B protein production promoting survival and cell movement in U2OS cells. |
| **IL4R** | **Exon 2 (SE), 5'UTR**  *Less exon inclusion upon eIF4E overexpression* | **7.47 ± 1.02**  **P < 0.005** | 2.31 ± 1.36  P = 0.22 | **3.82 ± 0.39**  **P < 0.005** | **3.79 ± 0.49**  **P < 0.005** | Exon 2 encodes 132 of 199 nt of the IL4R 5'UTR; reduced inclusion likely impacts translational regulation of the transcript. |
| **Shared targets (eIF4E + NCBP2)** | | | | | | |
| **MAPK8IP3** | **Exon 10 (SE)**  *Less included by both eIF4E and NCBP2* | **4.79 ± 0.71**  **P < 0.005** | **4.18 ± 0.81**  **P < 0.005** | **3.07 ± 0.53**  **P < 0.005** | **4.49 ± 0.68**  **P < 0.005** | Functional consequences of increased exon 10 inclusion are not yet known. |
| **QKI** | **Exon 3 (SE), KH domain**  *eIF4E decreased exon inclusion / NCBP2 increased exon inclusion* | **7.63 ± 4.55**  **P < 0.05** | **9.84 ± 2.06**  **P < 0.05** | **4.04 ± 0.96**  **P < 0.005** | **4.5 ± 1.49**  **P < 0.05** | Exon 3 encodes part of the KH domain used by all isoforms to bind RNA. Loss of this domain prevents QKI regulation of target mRNAs involved in differentiation, myelination, and blood vessel development. |
| **NCBP2-sensitive targets** | | | | | | |
| **UBAP2L** | **Exon 26a (SE)**  *Increased exon inclusion upon NCBP2 overexpression* | 1.63 ± 0.51  P = 0.16 | **4.12 ± 1.04**  **P < 0.05** | 2.06 ± 0.14  P = 0.12 | **3.38 ± 0.9**  **P < 0.05** | Exon 26a inclusion generates a minor isoform, UBAP2L-230. The functional consequence is unknown but could impact UBAP2L's role in stress granule formation. |
| **VEGFA** | **Exon 6 (SE), heparin-binding domain**  *Increased exon exclusion upon NCBP2 overexpression* | 1.80 ± 0.6  P = 0.13 | **6.10 ± 1.94**  **P < 0.05** | 1.77 ± 0.54  P =0.37 | **4.11 ± 1.56**  **P < 0.05** | Exon 7 encodes the majority of the heparin-binding domain of VEGFA. Increased inclusion allows more interactions with the extracellular matrix. |

*RT-qPCR and RNA immunoprecipitation (RIP) validation of eIF4E- and NCBP2-dependent alternative splicing events. Non-bold entries were non-significantly enriched in RIPs with the given cap-chaperone. We note that MAPK8IP3 does not appear in the ribodepletion list of alternatively spliced NCBP2 targets but is present in our list derived from poly-A isolation from the same cells (*[*GSE300496*](https://www.ncbi.nlm.nih.gov/geo/query/acc.cgi?acc=GSE300496) *versus* [*GSE295397*](https://www.ncbi.nlm.nih.gov/geo/query/acc.cgi?acc=GSE295397) *for polyA). This highlights the limitations of RNA-seq library preparation in fully capturing the targets.*
