## Supplementary material for "Nuclear RNA cap-chaperones eIF4E and NCBP2 influence distinct mRNA fates": Apendix 1

**Supplemental Tables**

**Table 1**. Metascape analysis of top 3 protein-protein interactions of differentially expressed transcripts following eIF4E overexpression

**Table 2**. Differentially expressed transcripts following NCBP2 overexpression in U2OS cells

**Table 3**. Gene Ontology analysis of differentially expressed transcripts following eIF4E overexpression

**Table 4**. Gene Ontology analysis of differentially expressed transcripts following NCBP2 overexpression

**Table 5**. Transcripts that are differentially expressed in both eIF4E and NCBP2 overexpression

**Table 6**. All significantly changing alternative splicing events following NCBP2 overexpression

**Table 7**. Significantly changing skipped exon and mutually exclusive exon events following NCBP2 overexpression

**Table 8**. Intron lengths surrounding skipped exon events following NCBP2 overexpression

**Table 9**. Transcripts with significant alternative splicing events in both eIF4E and NCBP2 overexpression

**Table 10**. Gene Ontology analysis of transcripts with alternative splicing events in both eIF4E and NCBP2 overexpression

**Table 11.** Gene Ontology analysis of transcripts alternatively spliced following NCBP2 overexpression

**Table 12**. Gene Ontology analysis of transcripts alternatively spliced following eIF4E overexpression

**Table 13**. Gene Ontology of transcripts with alternative spliced in mouse NCBP2 overexpression

**Table 14.** Skipped exon and mutually exclusive exon events in both eIF4E and NCBP2 overexpression

**Table 15.** Differentially expressed transcripts following eIF4E knockdown in U2OS cells

**Table 16**. Differentially expressed transcripts following NCBP2 knockdown in U2OS cells

**Table 17.** All significantly changing alternative splicing events following NCBP2 knockdown

**Table 18.** All significantly changing alternative splicing events following eIF4E knockdown

**Table 19.** Gene ontology analysis of transcripts with alternative splicing events in eIF4E knockdown

**Table 20.** Gene ontology analysis of transcripts with alternative splicing events in NCBP2 knockdown

**Table 21.** Transcripts with significant alternative splicing events in both eIF4E and NCBP2 knockdown

**Table 22.** Skipped exon and mutually exclusive exon events in both eIF4E and NCBP2 knockdown

**Table 23.** Gene Ontology analysis of transcripts with alternative splicing events in both eIF4E and NCBP2 knockdown

**Table 24.** Enriched transcripts in the FLAG-eIF4E pulldown vs input RIP-seq

**Table 25.** Gene Ontology analysis of transcripts enriched in FLAG-eIF4E RIP-seq

**Table 26.** Enriched transcripts in the FLAG-NCBP2 pulldown vs input RIP-seq

**Table 27.** Gene Ontology analysis of transcripts enriched in FLAG-eIF4E RIP-seq

**Table 28.** Transcripts enriched in FLAG-eIF4E and FLAG-NCBP2 RIP-seq

**Table 29**: Gene Ontology analysis of transcripts enriched in FLAG-eIF4E and FLAG-NCBP2 RIP-seq

**Table 30:** Antibodies used in western blots

**Table 31:** Primers used for qPCR
